# Fibroblast growth factor receptor substrate 2 interactome mapping reveals novel candidate interactors associated with migration and invasion

**DOI:** 10.1101/2025.09.23.678042

**Authors:** Levi Luca Kopp, Dejana Versamento, Bernard Ciraulo, Marc Thomas Schönholzer, Dina Hochuli, Meng-Syuan Lin, Martin Baumgartner

## Abstract

The scaffold protein FRS2 is central to FGFR signaling, linking receptor activation to MAPK/ERK and PI3K/AKT pathways. Elevated FRS2 expression correlates with aggressive tumor phenotypes and poor prognosis across multiple cancers, including the pediatric cerebellar tumor medulloblastoma (MB).

Here, we characterized FRS2’s subcellular localization and interactome in MB cells, employing live-cell imaging, phosphoproteomics, immunoprecipitation, and APEX2-based proximity labeling. We found that increased FRS2 expression is associated with increased motile and invasive behavior in MB tumor cells. We furthermore identified novel candidate FRS2-associated proteins involved in actin cytoskeleton remodeling, cell junction assembly, and translation initiation, which indicate a growth factor-dependent reorganization of the FRS2 signalosome. Our data furthermore indicate a regulatory role of FRS2 in directing subcellular distribution of the cell junction and cell motility regulator TJP1.

Our findings highlight the relevance of FRS2 as a mediator of cell motility and invasiveness and provide candidate proteins associated with FRS2 that are involved in cellular processes governing migration and invasion. This study thus provides a framework for exploring the FRS2 interactome as a possible target to attenuate FGFR-driven oncogenic processes with next-generation therapeutic strategies.

## Background

Signaling via receptor tyrosine kinases (RTKs) is relevant for the regulation of physiological processes including cell proliferation, differentiation and migration [1]. However, the dysregulation of RTK activity is also a major driver of pathological conditions, including cancer [1]. Fibroblast growth factor receptors (FGFRs) are a family of RTKs, frequently involved in oncogenic transformations [2]. Basic fibroblast growth factor (bFGF) ligand-induced pathway activation promotes local formation of cellular protrusions [3] and triggers the phosphorylation of proteins involved in cell adhesion and cell-cell interaction [4]. These observations imply that FGFR signaling regulates the reorganization of molecular structures, including the actin cytoskeleton and cell adhesion complexes in its vicinity. While the functional consequences of canonical FGFR pathway activation in health and disease are well explored, the molecular mechanism and critical interaction partners linking receptor activation to proximal regulators of biological processes and cytoskeleton remodeling remain poorly understood.

A central integrator of FGFR signaling is the fibroblast growth factor receptor substrate 2 (FRS2), a scaffold protein that coordinates the assembly of the intracellular receptor signalosome [5]. Its association with the plasma membrane is facilitated by a N-terminal myristoylation site [6], which is followed by a phosphotyrosine-binding (PTB) domain that constitutively binds a juxtamembrane region of FGFRs [7]. FRS2’s C-term is largely unstructured, harboring six tyrosine phosphorylation sites that serve as docking modules for adaptor proteins such as growth factor receptor-bound protein 2 (GRB2) or tyrosine-protein phosphatase non-receptor type 11 (SHP2) [6,8]. This modality suggests a binding-induced folding process of FRS2 that selectively connects FGFR activation to the downstream MAPK/ERK and PI3K/AKT signaling pathways.

FRS2 is a potential oncogene, whose overexpression has been reported in several cancers, including medulloblastoma [9], glioma [10], liposarcoma [11,12], as well as ovarian [13] and prostate cancers [14]. Further, elevated levels of FRS2 correlate with increased anchorage independent growth, proliferation and *in vivo* tumor formation [11,13]. Notably, this effect was stronger than constitutive activation of the FGFR pathway through MEK mutants, suggesting that FRS2 regulates additional pathways beyond canonical FGFR-MAPK signaling [13].

Low-throughput interaction studies identified few additional FRS2-binding partners potentially implicated in regulating proximal actin cytoskeleton structures and cell proliferation. These included Rho-related GTP-binding proteins RND1/2 [15], flotillin FLOT1 [16] and cyclin-dependent kinases regulatory subunits CKS1/2 [17]. FRS2 was detected in the proximal proteome of the actin regulators anillin (ANLN) [18], ezrin (EZR) [19] and myristoylated alanine-rich protein kinase C substrate (MARCKS) [19]. Collectively, these findings suggest that FRS2 exhibits broader functions beyond merely recruiting the canonical FGFR signaling pathway members to active receptors.

We hypothesize that in MB and other solid tumors, FRS2 may integrate FGFR signaling beyond the MAPK/ERK and PI3K/AKT pathways, into local networks of proteins involved in the regulation of cell growth, actin dynamics and cell motility. Given the key role of FRS2 in bFGF-induced invasion and tissue infiltration in MB [9], we specifically explored the interactome and proximal proteome of FRS2 in the context of growth factor-driven motile and invasive behavior of MB cells.

## Methods

### Primary tumor gene expression/alteration analysis

Pancancer analysis of FRS2 alterations and mRNA expression levels was performed on cBioPortal (https://www.cbioportal.org) using the TCGA PanCancer Atlas Studies collection (32 studies). A z-score threshold of -/+ 2, relative to average expression in diploid samples was used. Expression and survival data for MB patients was obtained from R2 genomics and visualization platform (https://r2.amc.nl). The following data sets were used: Berchtold (normal brain, 172, MAS5.0 – u133p2), Roth (normal cerebellum, 9, MAS5.0 – u133p2), Pfister (MB SHH, 73, MAS5.0 – u133p2), Hsieh (MB, 31, MAS5.0 – u133p2), Pfister (MB, 223, MAS5.0 – u133p2), Delattre (MB, 57, MAS5.0 – u133p2), Kool (MB, 62, MAS5.0 – u133p2), Gilbertson (MB, 76, MAS5.0 – u133p2) and Cavalli (MB, 763, rna_sketch–hugene11t), mixed Medulloblastoma / Cerebellum (re-analyses) – Swartling – 1641 – custom GSE124814 [20]. The GSE124814 dataset contains 1350 MB and 291 normal brain transcriptomic samples (cerebellar cortex, cerebellum, rhombic lip). Weishaupt *et al.* used a set of 100 genes with 25 signature genes to classify MB samples with available (1213) and samples without (137) subgroup affiliation. For MB subgroup and subtype analyses the Cavalli 2017 [21] nomenclature was applied. Kaplan–Meier survival curves were generated using the “scan” cutoff mode, which selects the most statistically significant expression cut point within the patient cohort.

Datasets were visualized in GraphPad Prism (v10.4.1) and differences in mRNA expression level Z-scores across multiple datasets were tested with ANOVA and Kruskal-Wallis tests. Different expression levels between two samples were tested with a Mann-Whitney test (unpaired, two-tailed). The statistical analyses are shown in Supplementary file 11.

### Cell lines and culture conditions

UW228 cells were generously provided by John Silber (Seattle, USA) and cultured in Dulbecco’s Modified Eagle Medium (DMEM, Sigma). D425-Med (D425) cells were kindly obtained by Henry Friedman (Durham, USA) and cultured in Minimum Essential Medium - Richter’s modification (IMEM, Sigma). ONS-76 cells were provided by Michael Taylor (Texas Children’s Hospital, USA) and cultured in RMPI-1640 medium (R0883, Sigma) supplemented with GlutaMAX (#35050-038, Gibco), 10% FBS (S0615, Sigma) and 1% penicillin/streptomycin (#15140-122, Gibco). HEK Lenti-X 293 were kindly provided by Beat Schäfer (University of Zürich, Switzerland) and cultured in Dulbecco’s Modified Eagle Medium (DMEM, Sigma). HAP1 wt (C631) and HAP1 FRS2 ^−^ (HZGHC007502c012) cells were purchased from Horizon Discovery (UK) and cultured in Iscove’s Modified Dulbecco’s Medium (IMDM, Gibco). For an overview of all cell lines, see Supplementary file 1. Cell line authentication and cross contamination testing was performed by Multiplexion by single nucleotide polymorphism (SNP) profiling. Cell lines were regularly tested for mycoplasma using PCR- or enzyme-based detection kits. Standard cell culture media were prepared by using 10% v/v heat-inactivated fetal bovine serum (FBS), 1% v/v Penicillin/Streptomycin, and 2 mM GlutaMAX (ThermoFisher). Serum-free media (SFM) is medium supplemented with 1% v/v P/S and 2 mM GlutaMAX. All tissue culture reagents were filtered using 0.22 µm syringe filters. The cells were maintained at 37 °C in a humidified atmosphere containing 5% CO_2_.

### Plasmids

Lentiviral constructs were ordered from VectorBuilder (Santa Clara, CA, USA). For the FRS2-mNeonGreen experiments, mNeonGreen was fused to the C-terminus of human FRS2 using a flexible 3xGS linker. The APEX2 sequence was adapted from Lam *et al.* [22] (Addgene plasmid #49386). For both plasmids, a blasticidin resistance gene was included for antibiotic selection. Vector details can be found in Supplementary file 1 and: https://en.vectorbuilder.com/vector/VB220316-1595dwh.html (FRS2-mNeonGreen) https://en.vectorbuilder.com/vector/VB221102-1154pbp.html (FRS2-APEX2-V5)

The lentiviral construct of LifeAct-EGFP (LA-EGFP) including an ampicillin resistance was kindly obtained from Oliver Pertz (University of Bern).

### Generation of FRS2-mNG/FRS2-APEX2-V5 overexpressing cell lines

mCherry-Nuc lentiviral particles were ordered from Takara (rLV.EF1.mCherry-Nuc-9, #0023VCT). All other lentiviral particles were produced by transfection of HEK Lenti-X 293 cells with 7.5 µg transfer plasmid, 4.5 µg psPAX2 (Addgene, #12260) and 3 µg pCMV-VSV-G plasmid (Addgene, #8454) using Lipofectamine 3000 transfection reagent (Invitrogen). Virus was harvested 30 hours post-transfection and filtered. UW228 cell lines were incubated with virus containing supernatant and 10 µg/mL polybrene for 48 hours. Filtered virus supernatants were stored at –80 °C until further use. Two days post-transduction, MB cells were washed and expanded. Antibiotic selection was started at day three post-transduction with 10 µg/mL blasticidin or 1 µg/mL puromycin dissolved in fresh complete medium. Selection was continued for two weeks, and the medium exchanged every 2-3 days. Protein expression was confirmed by immunoblotting and immunofluorescence assay.

For UW228 mCherry-Nuc, FRS2-mNeonGreen cells were further sorted by flow cytometry for positive mCherry-Nuc and high FRS2-mNG expression (top 5% of population).

### Immunoblotting (IB)

Cells were lysed using RIPA buffer supplemented with cOmplete Mini Protease inhibitor Cocktail (Roche, #11836153001) and phosphatase inhibitors PhosSTOP (Roche, #4906837001) and cleared by centrifugation (13’000 g, 10 min at 4 °C). Protein concentration was determined using the Pierce BCA Protein Assay Kit (ThermoFisher, #23225). Equal protein amount and concentrations were used for gel electrophoresis using 4-20% Mini-PROTEAN TGX Precast Protein Gels (Bio-Rad Laboratories AG, #4561094/4561096). Proteins were transferred to Trans-Blot Turbo 0.2 µm Nitrocellulose Transfer Membrane (Bio-Rad, #1704158/1704159) and membranes blocked with 5% w/v non-fat milk in TBST for 1 hour at RT. For antibodies used see Supplementary file 1. HRP-linked secondary antibodies were used with Western Blotting Substrate (ThermoFisher, #32209) or SuperSignal West Femto Maximum Sensitivity Substrate (ThermoFisher, #34095). Membranes were acquired using a ChemiDoc Imaging System (Bio-Rad, #12003153) and the integrated density of immune-reactive bands was quantified using ImageJ2 (v2.14.0/1.54f). For uncropped IB images see Supplementary file 10.

### Proliferation competition assay

UW228 wt and UW228 overexpressing FRS2-mNG/mCherry-Nuc were mixed in a 1:1 ratio and kept in culture for two weeks. Technical triplicates were prepared. Every 2-3 days cells were split and excess cells used to determine the FRS2-mNG/mCherry-Nuc positive population. Cells were washed with PBS and filtered into 5 mL polystyrene round-bottom tubes (Falcon/Brunschwig, #352235). Data was acquired using BD Fortessa Cell Analyzer and analyzed using FlowJo software (v10.8.0).

### Live cell tracking assay

A glass bottom 96-well plate (Cellvis, #P96-1.5H-N) was pre-coated with 50 µL 50 µg/mL PureCol bovine collagen (CellSystems, #5005) in water for one hour at RT. Plate was washed three times with 100 µL PBS and all liquid removed. 2’000 cells in 150 µL complete medium were seeded (UW228 mCherry-Nuc, UW228 LA-EGFP/mCherry-Nuc, UW228 FRS2-mNG/mCherry-Nuc low/high) and incubated overnight at 37 °C, 5% CO_2_. 50 µL complete medium ± 150 ng/mL bFGF was added to each well and incubated for 2 hours. Live imaging was performed using a Nikon Ti2 widefield microscope equipped with a climate chamber. Cells in each well were imaged for 15 h, whereby Cy3 channel images were acquired every 3 min using a 10x objective (0.73 µm/px). 12 wells were acquired per condition.

Live-cell imaging data was analyzed using Fiji’s TrackMate plugin. Nuclei were detected using LogDetectorFactory with enabled subpixel localization. Tracking was conducted with Sparse LAP Tracker using a maximum linking distance of 50 px. Gap closing, track splitting and track merging were disabled. Single cell trajectory features were calculated and exported including mean speed, displacement and duration. Directionality was calculated dividing displacement by total distance traveled (mean speed x track duration). Tracks were filtered for a minimal duration of 30 frames. Further, tracks with zero or negative mean speed, zero or negative displacement, and directionality values bigger than one were removed. About 80% of all tracks passed these filtering criteria. Mean track speed and directionality were plotted as a function of cell line and growth factor presence.

Statistical significance between the track mean speed values were calculated using a Student’s t test (unpaired, two-tailed). For the directionality data set a Mann Whitney U test was applied.

### Spheroid invasion assay (SIA)

SIA was performed as described [23]. In brief: 2’500 cells were seeded per well in 100 µL complete medium in a 96-well clear round bottom ultra-low mount microplate (Corning Costar, #7007) and incubated overnight at 37 °C, 5% CO_2_. After 24 hours, the formation of compact and uniform spheroids was confirmed by light microcopy. 70 µL medium was removed and replaced with 70 µL PureCol bovine collagen (2.7 mg/mL, CellSystems, #5005). Polymerized collagen hydrogels were overlaid with 100 µL SFM with optional addition of recombinant human FGF-basic (final concentration 100 ng/mL, Peprotech, #100-18B). The cells were allowed to invade the collagen matrix for 24 hours, after which they were stained with Hoechst (1:2000, Sigma Aldrich, #B2883) for 3-4 hours. Image acquisition was performed with an Operetta CLS High-Content Analysis System (PerkinElmer, #HH16000000) using the 405 nm channel. Spheroids and invading cells were defined based on fluorescence threshold using Harmony software. For each tumor cell nucleus, the distance from the center of the spheroid was calculated. The total distance of invasion from the center of the spheroid was calculated by summing up the individual invasion distances of each cell using Harmony. The data was displayed in GraphPad Prism (v10.4.1). Statistical analysis was performed using an ANOVA with multiple comparison tests.

### Live imaging of FRS2-mNG

200 µL complete medium with 5x10^3^ UW228 FRS2-mNG/mCherry-Nuc cells were seeded per well in an Ibidi µ-slide 8-well collagen IV plate (Ibidi, #80822) and incubated for 6 hours at 37 °C, 5% CO_2_. Live-cell imaging was performed using a Nikon Ti2 widefield microscope equipped with a climate chamber. Cells were imaged for 16 hours with pictures acquired every 3 min by using a 40x objective (2424 x 2424 px, 0.18 µm/px) in the GFP and Cy3 channel. Using ImageJ2 (v2.14.0/1.54f) we adjusted brightness and contrast and added time stamp and scale bars.

### Immunofluorescence assay (IFA)

Coverslips were coated with collagen I in water (PureCol (CellSystems, #5005), 50 µg/mL final, 0.5 mL/well of 24 well plate,) for 1 hour at RT and then washed with PBS. 500 µL complete medium with 1.5x10^4^ UW228 FRS2-mNG cells were seeded per well and incubated overnight. Cells were fixed in complete medium by adding PFA (to 4% v/v) for 20 min at 37 °C. Cells were washed 2 x 5 min in PBS, permeabilized with 0.1% v/v Triton X-100 in PBS for 20 min at RT and washed again 3 x 5 min in PBS. Cells were blocked with PBS supplemented with 5% v/v FBS for 1 hour at RT. For a list of the used primary and secondary antibodies, see Supplementary file 1. Primary antibodies were diluted in blocking buffer (PBS, 5% v/v FBS) and incubated overnight at 4 °C in a humified chamber. Cells were washed 3 x 5 min in PBS and incubated with secondary antibodies for 1 hour at RT. Cells were washed 3 x 5 min in PBS and nucleic acid stained with DAPI (1:5000) diluted in PBS for 5 min at RT. Coverslips were mounted using DAKO Glycergel mounting medium (DAKO, #C0563). Image acquisition was performed on a NikonTi2 widefield microscope.

### RNA interference

ONS-76 or UW228 cells at approximately 70% confluency were transfected with Lipofectamine RNAiMAX Transfection Reagent (13778075, Thermo Fisher Scientific) and 10 – 20 nM siRNAs following the manufacturer’s instruction. siRNAs used are listed in below. Non-targeting control siRNA was used as a negative control. After 48 h, RNA and proteins were isolated from the cells to determine gene expression by RT-qPCR and to evaluate protein expression by IB. For the SIA, cells were re-seeded to form spheroids in ultra-low adhesion plates 24 h after the transfection. For the live cell imaging and immunofluorescence assays, cells were re-seeded after 24 h after the transfection. siRNA sequences can are listed in Supplementary file 1.

### RT-qPCR analysis of gene expression

125’000 HAP1 cells/well were seeded in 6-well plates and incubated at 37 °C overnight. RNA was extracted and purified using the RNeasy Plus Mini kit (Qiagen, 74134) according to the manufacturer’s protocol and stored at –80 °C. RNA concentration was measured using a Nanodrop instrument. High-Capacity cDNA Reverse Transcription kit (ThermoFisher 4374966) was used to prepare cDNA according manufacturer’s protocol. A total amount of 1 µg RNA was diluted in 10 µL RNAse-free water and mixed with 10 µL master mix, containing RT buffer, dNTPs, RT random primers and MultiScribe Reverse transcriptase before performing reverse transcription in a thermocycler.

Per sample, 9 µL TaqMan Universal PCR Master Mix (ThermoFisher, 4444556) containing the specific primers (see Supplementary file 1) was prepared and mixed with 1 µL cDNA. Triplicates for each sample were prepared in a 384-well plate. The plate was sealed with a MicroAmp Optical Adhesive film (ThermoFisher, 4311971) and centrifuged for 1 min at 800 rpm. The RT-qPCR was performed using a 7900HT Fast Real-Time PCR system and gene expression calculated according the 2^−ΔΔCT^ method [24]. Expression of target genes were normalized to GAPDH and the data visualized using GraphPad Prism (v10.4.1). Statistical analysis was performed using a Student’s t test (unpaired, two-tailed).

### Phospho-proteomics assay

#### Sample preparation

6-well plates were seeded with 3x10^5^ D425 wt cells/well in complete medium in cell culture 6-well plates. The next day, the medium was replaced with serum-free medium and the cells starved overnight. Three biological replicates (n=3) were prepared for each treatment condition (PBS and bFGF). Cells were pre-treated with 0.1% v/v DMSO for 1 h, followed by stimulation with either 50 ng/mL bFGF or PBS for 5 min. Plates were placed on ice immediately after bFGF stimulation, the medium was removed and the cells lysed in 150 µL high-SDS RIPA containing 4% w/v SDS, 1x cOmplete protease inhibitors, 1x Halt Phosphatase Inhibitor Single-Use Cocktail (ThermoFisher 78428). Lysates were transferred to pre-cooled Eppendorf tubes and snap-froze in liquid nitrogen. Samples were stored at –80 °C for later processing.

#### Sample digestion and clean up

A volume corresponding to 150 µg was taken from each sample and treated with High Intensity Focused Ultrasound (HIFU) for 1 minute at an ultrasonic amplitude of 100%. The samples were boiled at 95 °C for 10 min while shaking at 800 rpm on a Thermoshaker (Eppendorf) and subsequently centrifuged at 20’000 g for 10 min. Sample digestion was carried out using a modified on-filter digestion protocol of the filter-aided sample preparation (FASP) protocol [25]. Briefly, proteins were diluted in 200 µL of UT buffer (Urea 8 M in 100 mM Tris/HCL pH 8.2), loaded on Ultracel 30000 MWCO centrifugal unit (Amicon Ultra, Merck, Darmstadt, Germany) and centrifuged at 14’000 g. SDS buffer was exchanged by one centrifugation round of 200 µL UT buffer. Proteins were reduced with 2 mM TCEP (tris(2-carboxyethyl)phosphine) and alkylated with 15 mM chloroacetamide at 30 °C for 30 min, followed by three 100 µL washing steps with UT and two 100 µL washing steps with Triethylammonium bicarbonate buffer (TEAB, pH 8). Finally, proteins were on-filter digested using 120 µL of 0.05 TEAB (pH 8) containing trypsin (Promega, Madison, WI, USA) in ratio 1:50 (w/w). Digestion was performed overnight in a wet chamber at room temperature, and peptides were eluted by centrifugation at 14’000 g for 25 minutes. Peptide concentration was determined using the Lunatic UV/Vis polychromatic spectrophotometer (Unchained Labs) before drying the samples to completeness.

#### TMT labelling and peptide fractionation

250 µg TMTpro 18-plex reagent (Thermo Fisher Scientific) was dissolved in 15 μl of anhydrous acetonitrile (Sigma-Aldrich) and added to 75 µg peptides in 45 µL of 50 mM TEAB, pH 8.5. The solution was gently mixed and incubated for 60 min at RT. The reaction was quenched by adding 8 µL of 5% hydroxylamine (Thermo Fisher Scientific). The combined TMT sample was created by mixing equal amounts of each TMT channel together. Labeled peptides were offline pre-fractionated using high pH reverse phase chromatography. Peptides were separated on an XBridge Peptide BEH C18 column (130Å, 3.5 µm, 4.6 mm x 250 mm, Waters) using a 72 min linear gradient from 5-40% acetonitrile/9 mM NH_4_HCO_2_. Every minute a new fraction was collected and concatenated into 12 final fractions.

#### Phosphopeptide enrichment

The phosphopeptide enrichment was performed using a KingFisher Flex System (Thermo Fisher Scientific) and PureCube Fe-NTA MagBeads (Cube Biotech). Beads were conditioned following the manufacturer’s instructions, consisting of 3 washes with 200 µL of binding buffer (80% acetonitrile, 0.1% TFA). Each fraction was dissolved in 200 µL binding buffer. The beads, wash solutions and samples were loaded into 96-well microplates and transferred to the KingFisher. Phosphopeptide enrichment was carried out using the following steps: washing of the magnetic beads in binding buffer (5 min), binding of the phosphopeptides to the beads (30 min), washing the beads in wash 1-3 (binding buffer, 3 min each) and eluting peptides from the beads (50 µL 2.5% NH_4_OH in 50% acetonitrile, 10 min). Each elution was combined with 30 µL neutralisation solution (75% acetonitrile, 10% formic acid) and samples were dried to completeness and re-solubilized in 10 µL of 3% acetonitrile, 0.1% formic acid for MS analysis. For pTyr enrichment, 12 µg pTyr1000 and 12 µg 4G10 anti-pTyr antibodies were immobilized on Protein G agarose beads in IP buffer (100mM Tris-HCl/0.4% NP-40) and incubated with the TMT-labeled peptide pool rotating overnight at 4 °C. Beads were washed with IP buffer and deionized water before elution in FeNTA binding buffer (Thermo). Peptides eluted from the antibody mix were incubated with FeNTA spin columns to further separate phosphorylated peptides from unspecific background binders. After elution, the peptides were dried to near-completeness and transferred to Evotips.

#### LC-MS/MS analysis

Mass spectrometry analysis was performed on an Orbitrap Exploris 480 mass spectrometer (Thermo Fisher Scientific) equipped with a Nanospray Flex Ion Source (Thermo Fisher Scientific) and coupled to an M-Class UPLC (Waters). Solvent composition at the two channels was 0.1% formic acid for channel A and 0.1% formic acid, 99.9% acetonitrile for channel B. Column temperature was set to 50 °C. For phosphopeptide analysis, 4 µL were injected, while full proteome analysis was carried out using a portion of each fraction before phosphopeptide enrichment. Peptides were loaded on a commercial nanoEase MZ Symmetry C18 Trap Column (100Å, 5 µm, 180 µm x 20 mm, Waters) connected to a nanoEase MZ C18 HSS T3 Column (100Å, 1.8 µm, 75 µm x 250 mm, Waters). Peptides were eluted at a flow rate of 300 nL/min. After a 3 min initial hold at 5% B, a gradient from 5 to 22% B in 80 min and 22 to 32% B in additional 10 min was applied. The column was cleaned after the run by increasing to 95% B and holding 95% B for 10 min prior to re-establishing loading condition for another 10 minutes. Peptides loaded onto Evotips were analyzed on an Evosep One system using the 20 SPD Whisper method with an Aurora Elite column connected.

The mass spectrometer was operated in data-dependent mode (DDA) with a maximum cycle time of 3 s, with spray voltage set to 2.4 kV, funnel RF level at 40% and heated capillary temperature at 275 °C. Full-scan MS spectra (350−1’500 m/z) were acquired at a resolution of 120’000 at 200 m/z after accumulation to a target value of 3’000’000 or for a maximum injection time of 45 ms. Precursors with an intensity above 5’000 were selected for MS/MS. Ions were isolated using a quadrupole mass filter with 0.7 m/z isolation window and fragmented by higher-energy collisional dissociation (HCD) using a normalized collision energy of 32%. HCD spectra were acquired at a resolution of 60’000 (phosphopeptides) or 30’000 with turboTMT on (full proteome) and maximum injection time was set to 200 ms (phosphopeptides) or Auto (full proteome). The normalized automatic gain control (AGC) was set to 100%. Charge state screening was enabled such that singly, unassigned and charge states higher than six were rejected. Precursor masses previously selected for MS/MS measurement were excluded from further selection for 20 s, and the exclusion window was set at 10 ppm. The samples were acquired using internal lock mass calibration on m/z 371.1012 and 445.1200.

The mass spectrometry proteomics data were handled using the local laboratory information management system (LIMS) [26] and all relevant data have been deposited to the ProteomeXchange Consortium via the PRIDE (http://www.ebi.ac.uk/pride) partner repository with the data set identifier PXD062602.

#### Data analysis

Thermo .raw files were processed with Proteome Discoverer 2.5 and searched against the reviewed human reference proteome database UP000005640 (download from 2023-01-27) appended with common contaminants using Sequest and Percolator for FDR calculation (0.01 filter on PSM and peptide level). Determination of site localization probability was done using the IMP-ptmRS node with default cut-off filter of ≥0.75. Full proteome and phospho-enriched samples were processed separately. For statistical evaluation and site-specific integration of results, the peptideGroups.txt output of Proteome Discoverer for phospho-enriched datasets were filtered for only confidently assigned phosphorylated peptides, and for each phospho-peptide assignment the number of phosphorylations, the modified residue and position in protein were parsed into a data frame. Further, a new identifier was generated comprising of protein accession, the phospho-acceptor residue and the phospho-site in the full-length protein. This identifier was then used to summarize (sum) quantities from each TMT-channel. This site-centric data is then further analyzed using the prolfqua R-package [27].

For this, we first filter for a minimum abundance of 1 for each TMT-channel abundance, then the data is log2-transformed and normalized with a robust z-score transformation. A linear model is fitted to each protein-phosphosite abundance to compute significant differences between different conditions. Group comparisons (bFGF_DMSO vs. SFM_DMSO) were evaluated with a moderated Wald-test with pooled variance (as implemented in the limma R-package [28]). The resulting p-values were adjusted for multiple testing using BH. In a very similar way, the rolled-up to protein results from Proteome Discoverer was used and the same data analysis steps were applied as for the phospho-peptide results. After assessing differential expression on protein level using prolfqua these protein centric results where then joined with the phosphosite-centric results.

### Co-immuno-precipitation (Co-IP)

2 x 10^6^ UW228 FRS2-mNG/mCherry-Nuc and UW228 WT control cells were seeded in 20 mL complete medium in 15 cm cell culture dishes and incubated overnight at 37 °C, 5% CO_2_. The next day, medium was replaced with SFM and cells starved overnight. bFGF was added to a final concentration of 50 ng/mL and cells were incubated for 5 min at 37 °C. Medium was removed, 200 µL cold lysis buffer (10 mM Tris, 150 mM NaCl, 0.5 mM EDTA, 0.25% v/v NP-40, pH 7.5) was added, and cells were removed by scraping. Cells were lysed for 10 min on ice and lysates cleared by centrifugation (10 min, 13’000 g, 4 °C). Protein concentration was determined with Pierce BCA Protein Assay Kit (ThermoFisher, #23225). For all samples, 200 µL lysate with equal concentrations was prepared and diluted with 300 µL cold dilution buffer (10 mM Tris, 150 mM NaCl, 0.5 mM EDTA, pH 7.5). Per sample, 25 µL mNeonGreen-Trap Magnetic Agarose beads (Chromotek, NTMA-10) were prepared by washing 3x with 500 µL dilution buffer. The lysates of UW228 FRS2-mNG/mCherry-Nuc and of the UW228 WT control cells were mixed with the beads and incubated for 90 min at 4 °C. Supernatant was removed and beads were washed 3x with 500 µL cold wash buffer (10 mM Tris, 150 mM NaCl, 0.5 mM EDTA, 0.05% v/v NP-40, pH 7.5).

Alternatively, for pull-down assays using the FGFR-binding peptide: Per sample 35 µL Dynabeads MyOne Streptavidin T1 (Invitrogen, #65601) were washed twice in 500 µL PBST (0.02% v/v Tween-20) and resuspended in 100 µL PBST. Beads were coated with 20 µL 2 nmol/µL biotinylated FGFR peptide (LifeTein, biotin-GGGGSHSQMAVHKLAKSIPLRRQVTVS) for 1 hour at RT. Beads were washed twice with 500 µL before incubating them with the lysates for 90 min at 4 °C and 90 min at RT. Beads were washed twice with 500 µL lysis buffer.

For immunoblot analysis, beads were transferred to new Eppendorf tubes and eluted by boiling for 5 min at 95 °C in 60 µL elution buffer (40 µL wash buffer, 20 µL 4 x Laemmli Sample Buffer (Bio-Rad, #1610747)). 25% of the eluate was used per immunoblot to detect co-precipitated proteins. For mass spectrometry analysis of the mNeonGreen IPs, beads were transferred to new Eppendorf tubes with the last washing step and kept on ice till further processing for MS.

#### Sample digestion and clean up

For each sample (*n*=3), the washed beads were resuspended in 45 µL digestion buffer (triethylammonium bicarbonate (TEAB), pH 8.2), reduced with 5 mM TCEP (tris(2-carboxyethyl)phosphine) and alkylated with 15 mM chloroacetamide. Proteins were on-bead digested using 500 ng of Sequencing Grade Trypsin (Promega). The digestion was carried out at 37 °C overnight. The supernatants were transferred into new tubes and the beads were washed with 150 µL trifluoroacetic acid (TFA) -buffer (0.1% TFA, 50% acetonitrile) and combined with the first supernatant. The samples were dried to completeness and re-solubilized in 20 µL of MS sample buffer (3% acetonitrile, 0.1% formic acid). Peptide concentration was determined using the Lunatic UV/Vis polychromatic spectrophotometer (Unchained Labs).

#### LC-MS/MS data acquisition

Mass spectrometry analysis was performed on an Orbitrap Exploris 480 mass spectrometer (Thermo Fisher Scientific) equipped with a Nanospray Flex Ion Source (Thermo Fisher Scientific) and coupled to an M-Class UPLC (Waters). MS solvent composition at the two channels was 0.1% formic acid for channel A and 0.1% formic acid, 99.9% acetonitrile for channel B. Column temperature was set to 50 °C. For each sample, peptides corresponding to 0.2 absorbance measurement were loaded on a commercial nanoEase MZ Symmetry C18 Trap Column (100Å, 5 µm, 180 µm x 20 mm, Waters) in-line connected to a nanoEase MZ C18 HSS T3 Column (100Å, 1.8 µm, 75 µm x 250 mm, Waters). The peptides were eluted at a flow rate of 300 nL/min with a gradient from 5 to 22% B in 40 min and 22 to 32% B in an additional 5 min. The column was cleaned after each run by increasing to 95% B and prior to re-establishing loading condition for another 10 minutes.

Peptides from AP-MS experiments were acquired in data-independent mode (DIA). DIA scans covered a range from 350 to 1050 m/z in windows of 10 m/z. The resolution of the DIA windows was set to 30’000, with a normalized AGC target value of 3’000%, the maximum injection time set to Auto and a fixed normalized collision energy (NCE) of 28%. Each instrument cycle was completed by a full MS scan monitoring 350 to 1’050 m/z at a resolution of 120’000. The samples were acquired using internal lock mass calibration on m/z 371.1012 and 445.1200.

The mass spectrometry proteomics data were handled using the local laboratory information management system (LIMS) [26] and all relevant data have been deposited to the ProteomeXchange Consortium via the PRIDE (http://www.ebi.ac.uk/pride) partner repository with the data set identifier PXD062611.

#### LC-MS/MS data analysis

The acquired DIA data were processed for identification and quantification using DIA-NN 1.8.2 [29]. Spectra were searched against the Uniprot Homo sapiens reference proteome (UP000005640, downloaded 2023-08-24), concatenated to its reversed decoyed FASTA database and common protein contaminants. Carbamidomethylation of cysteine was set as a fixed and methionine oxidation set as variable modification. Enzyme specificity was set to trypsin/P, allowing a minimal peptide length of 7 amino acids and a maximum of one missed cleavage.

Interactor scoring from three biological replicates of UW228 FRS2-mNG/mCherry-Nuc was done using the SAINTexpress algorithm [30] applied to intensities as protein abundance estimates. In short, a set of R functions implemented in the R package prolfqua was used to convert the intensity protein abundances reported in the combined_protein.txt file into SAINTexpress-compatible format. The interactor-scoring (enrichment vs background) was done by comparison of the UW228 FRS2-mNG to the UW228 WT pull-down. A comparable number of proteins detected by a minimum of 2 peptides per protein across all conditions was recorded (UW228 FRS2-mNG: 3338, 3910 and 3941; UW228 wt: 3691, 3791, 3805).

Obtained data sets were screened for significant hits by filtering enriched proteins with a log_2_FC ≥ 1 and bFDR ≤ 0.05. Hits with a bFDR of 0 were adjusted to 0.0001 to enable plotting of highly significant interactors.

### APEX2-mediated proximity labeling assay

#### Sample preparation

The protocol for AEPX2-based proximity biotinylation was adapted from Tan *et al.* 2020, STAR Protocols [31]. The FRS2-APEX2-V5 was used as bait and FRS2-mNG included as a no-APEX negative control (background) treated identically as the FRS2-APEX. The interactor-scoring (enrichment vs background) was done by comparison of the FRS2-APEX to the no-APEX control. A no-APEX control was chosen because omission of APEX itself yields the lowest background as endogenous peroxidase activity can lead to non-specific labeling and false positive results. A free-floating (cytosolic) APEX would cause substantial background biotinylation. Three biological replicates were generated per condition. For each replicate, two 15 cm cell culture dishes were seeded with 20 mL complete medium containing 2.1 x 10^6^ cells (UW228 FRS2-APEX2-V5 or UW228 FRS2-mNG/mCherry-Nuc) and incubated overnight. The next day, the medium was replaced with 15 mL SFM per dish and cells were starved overnight. On day 3, SFM was reduced to 10 mL per dish, 10 µL DMSO were added per plate and cells were Incubated for 30 min at 37 °C. Next, 101 µL 100x biotin phenol stock (final concentration: 2.5 mM) in DMSO was added and mixed until completely dissolved. Cells were incubated another 30 min at 37 °C. For the last 3 min 1 mL of SFM only or SFM + bFGF (final concentration 50 ng/mL) was added, and immediately afterwards 1 mL 6 mM H_2_O_2_ in PBS (final H_2_O_2_ concentration: 0.5 mM). The stimulation/biotinylation reaction was performed for 3 min at 37 °C and then quenched by the addition of 12 mL 2x STOP buffer (1 mM MgCl_2_, 2 mM CaCl_2_, 10 mM Trolox, 20 mM sodium ascorbate, 20 mM sodium azide, PBS). Thorough mixing was achieved by slightly swirling the medium for 1 min. After removal of the reaction solution, the cells were washed twice with 10 mL 1x STOP buffer (0.5 mM MgCl_2_, 1 mM CaCl_2_, 5 mM Trolox, 10 mM sodium ascorbate, 10 mM sodium azide, PBS) and once with 15 mL PBS. 4 mL trypsin/EDTA was then added for 3 min at 37 °C to detach cells. Cells were collected in 12 mL complete medium and identical conditions were pooled before centrifugation for 5 min, 300 g at RT. The cell pellet was washed with 12.5 mL PBS and centrifuged for 5 min, 300 g, at RT.

After PBS removal cells were lysed in 2 mL lysis buffer (5 mM Trolox, 10 mM sodium ascorbate, 10 mM sodium azide, RIPA) containing cOmplete Mini Protease inhibitor Cocktail (Roche, #11836153001) and phosphatase inhibitors PhosSTOP (Roche, #4906837001) per sample. Cells were incubated for 30 min on ice before each sample was sonicated twice for 30 sec, 10% power. Lysates were cleared by centrifugation (20 min, 13’000 g, 4 °C) and proteins precipitated by addition of 5-fold volume cold acetone (–20 °C). Samples were vortexed and incubated for 3 hours at –20 °C before centrifugation for 15 min, 2’500 g, at RT. Acetone supernatant was discarded and protein pellet completely air-dried for 1 min. 2 mL RIPA buffer were added (incl. inhibitors) and protein pellets were redissolved. Residual pellet pieces were resolubilized by using a Douncer lyser. Samples were sonicated twice for 30 sec (10% power) and cleared by centrifugation (15 min, 13’000 g, 4 °C). Lysates were snap-frozen in liquid nitrogen and stored at –80 °C.

After the collection of all samples, lysates were thawed on ice and protein concentrations measured by using Pierce BCA Protein Assay Kit (ThermoFisher, #23225). 6 mg total protein was prepared for each sample in 1.8 mL RIPA buffer (incl. inhibitors). Per sample, 30 µL streptavidin-coated magnetic beads (Dynabeads™ MyOne™ Streptavidin T1, Invitrogen, #65601) were washed twice in RIPA before being incubated with lysates overnight at 4 °C.

Beads were washed twice with 1 mL RIPA, once with 1 mL RIPA high SDS (1% w/v SDS), once with 1 mL RIPA, and finally with 1 mL TBS (20 mM Tris, 150 mM NaCl, pH 7.6). Beads in solution were transferred to new Eppendorf tube, washed with 100 µL TBS and transferred a second time to a new Eppendorf tube. Tubes were kept on ice until further processing.

#### Sample digestion and clean up

For each sample, the washed beads were resuspended in 45 µL digestion buffer (triethylammonium bicarbonate (TEAB), pH 8.2), reduced with 5 mM TCEP(tris(2-carboxyethyl)phosphine) and alkylated with 15 mM chloroacetamide. Proteins were on-bead digested using 500 ng of Sequencing Grade Trypsin (Promega). The digestion was carried out at 37 °C overnight. The supernatants were transferred into new tubes and the beads were washed with 150 µL trifluoroacetic acid (TFA) -buffer (0.1% TFA, 50% acetonitrile) and combined with the first supernatant. The samples were dried to completeness and re-solubilized in 20 µL of MS sample buffer (3% acetonitrile, 0.1% formic acid). Peptide concentration was determined using the Lunatic UV/Vis polychromatic spectrophotometer (Unchained Labs).

#### LC-MS/MS data acquisition

Mass spectrometry analysis was performed on an Orbitrap Exploris 480 mass spectrometer (Thermo Fisher Scientific) equipped with a Nanospray Flex Ion Source (Thermo Fisher Scientific) and coupled to an M-Class UPLC (Waters). MS solvent composition at the two channels was 0.1% formic acid for channel A and 0.1% formic acid, 99.9% acetonitrile for channel B. Column temperature was set to 50 °C. For each sample, peptides corresponding to 0.2 absorbance measurement were loaded on a commercial nanoEase MZ Symmetry C18 Trap Column (100Å, 5 µm, 180 µm x 20 mm, Waters) in-line connected to a nanoEase MZ C18 HSS T3 Column (100Å, 1.8 µm, 75 µm x 250 mm, Waters). The peptides were eluted at a flow rate of 300 nL/min with a gradient from 5 to 22% B in 40 min and 22 to 32% B in an additional 5 min. The column was cleaned after each run by increasing to 95% B and prior to re-establishing loading condition for another 10 minutes.

Peptides from proximity labeling experiments were analysed in data-dependent mode (DDA) with a maximum cycle time of 3 s, using Xcalibur (tune version 4.1 (DDA) or 4.3 (DIA)), with spray voltage set to 2.5 kV, funnel RF level at 40%, heated capillary temperature at 275 °C, and Advanced Peak Determination (APD) on. Full-scan MS spectra (350−1’200 m/z) were acquired at a resolution of 120’000 at 200 m/z after accumulation to a target value of 3’000’000 or for a maximum injection time of 45 ms. Precursors with an intensity above 5’000 were selected for MS/MS. Ions were isolated using a quadrupole mass filter with 1.2 m/z isolation window and fragmented by higher-energy collisional dissociation (HCD) using a normalized collision energy of 30%. HCD spectra were acquired at a resolution of 30’000 and maximum injection time was set to 119 ms. The automatic gain control (AGC) was set to 100’000 ions. Charge state screening was enabled such that singly, unassigned and charge states higher than seven were rejected. Precursor masses previously selected for MS/MS measurement were excluded from further selection for 20 s, and the exclusion window was set at 10 ppm.

The mass spectrometry proteomics data were handled using the local laboratory information management system (LIMS) [26] and all relevant data have been deposited to the ProteomeXchange Consortium via the PRIDE (http://www.ebi.ac.uk/pride) partner repository with the data set identifier PXD062524.

#### LC-MS/MS data analysis

The acquired DIA data were processed for identification and quantification using DIA-NN 1.8.2 [29].Spectra were searched against the Uniprot Homo sapiens reference proteome (UP000005640, downloaded 2023-08-24), concatenated to its reversed decoyed FASTA database and common protein contaminants. Carbamidomethylation of cysteine was set as a fixed and methionine oxidation set as variable modification. Enzyme specificity was set to trypsin/P, allowing a minimal peptide length of 7 amino acids and a maximum of one missed cleavage.

Interactor scoring was done using the SAINTexpress algorithm [30] applied to intensities as protein abundance estimates. In short, a set of R functions implemented in the R package prolfqua was used to convert the intensity protein abundances into SAINTexpress-compatible format. A comparable number of proteins detected by a minimum of 2 peptides per protein was recorded (UW228 FRS2-APEX + bFGF: 1766, 1759 and 1749, UW228 FRS2-APEX no bFGF: 1762, 1762 and 1760). Obtained data sets were screened for significant hits by filtering enriched proteins with a log_2_FC ≥ 1 and FDR ≤ 0.05. Minimum FDR values were adjusted to 0.0001 to enable plotting of highly significant hits with and FDR of 0.

### Downstream processing of protein data sets

#### Over-representation analysis (ORA)

ORA was performed using Web-based Gene Set Analysis Toolkit WebGestalt (https://www.webgestalt.org). Significantly enriched proteins from FRS2-mNG immunoprecipitation or APEX2 labeling assays (FDR ≤ 0.05, log_2_FC ≥ 1) where screened for enriched gene ontology (biological process noRedundant) terms. The built-in genome protein-coding data set was selected as reference. Enriched gene sets were extracted and plotted using GraphPad Prism (v10.4.1).

#### Protein network analysis

Protein network analysis was performed using the STRING database (https://string-db.org). Significant proteins (FDR ≤ 0.05, Log2FC ≥ 1) from the respective data sets were used to predict a protein interaction network. Text mining, experiments and databases were used as active interaction sources and minimum required interaction score was set to high confidence (0.700).

## Results

### Increased expression of FRS2 correlates with aggressive subtypes and worse outcomes in multiple cancers

The scaffold protein FRS2 plays a crucial role in the signalosome assembly of active FGF-receptors [5–8,32,33]. We previously reported that FRS2 function is essential for bFGF-driven migration and invasion in medulloblastoma (MB) [9,23,34,35]. To explore the potential clinical relevance of FRS2 expression across cancer types, we analyzed mRNA levels and genomic alterations using the publicly available cancer genomic data from the TCGA PanCancer atlas and the R2: Genomics Analysis and Visualization Platform.

Elevated *FRS2* mRNA levels (z-score > 2, relative to average expression in diploid samples) were detected in 7% of the 10’953 PanCancer patients. Patients with lung, esophagogastric, bladder, breast, sarcoma and head/neck cancer comprised the majority of all cases with dysregulated *FRS2* levels (Figure 1A). Normalized to their occurrence, sarcoma and bladder cancer patients showed particularly high incidences of elevated *FRS2* levels in 15-22% of all patients (Figure 1B). In the PanCancer patient dataset, elevated *FRS2* mRNA levels were associated with a negative impact on overall survival (Figure S1A). Conversely, while the mRNA levels of the *FRS2* homolog *FRS3* were also increased in 7% of patients (Figure S1B,C), no change in the overall survival was observed (Figure S1D). At genomic level, *FRS2* alterations were detected with an incidence of 3% of all tested patients. Gene amplifications, being the most frequent (70%), could explain increased mRNA levels in some of the patients (Figure 1C, S1E).

**Figure 1.**
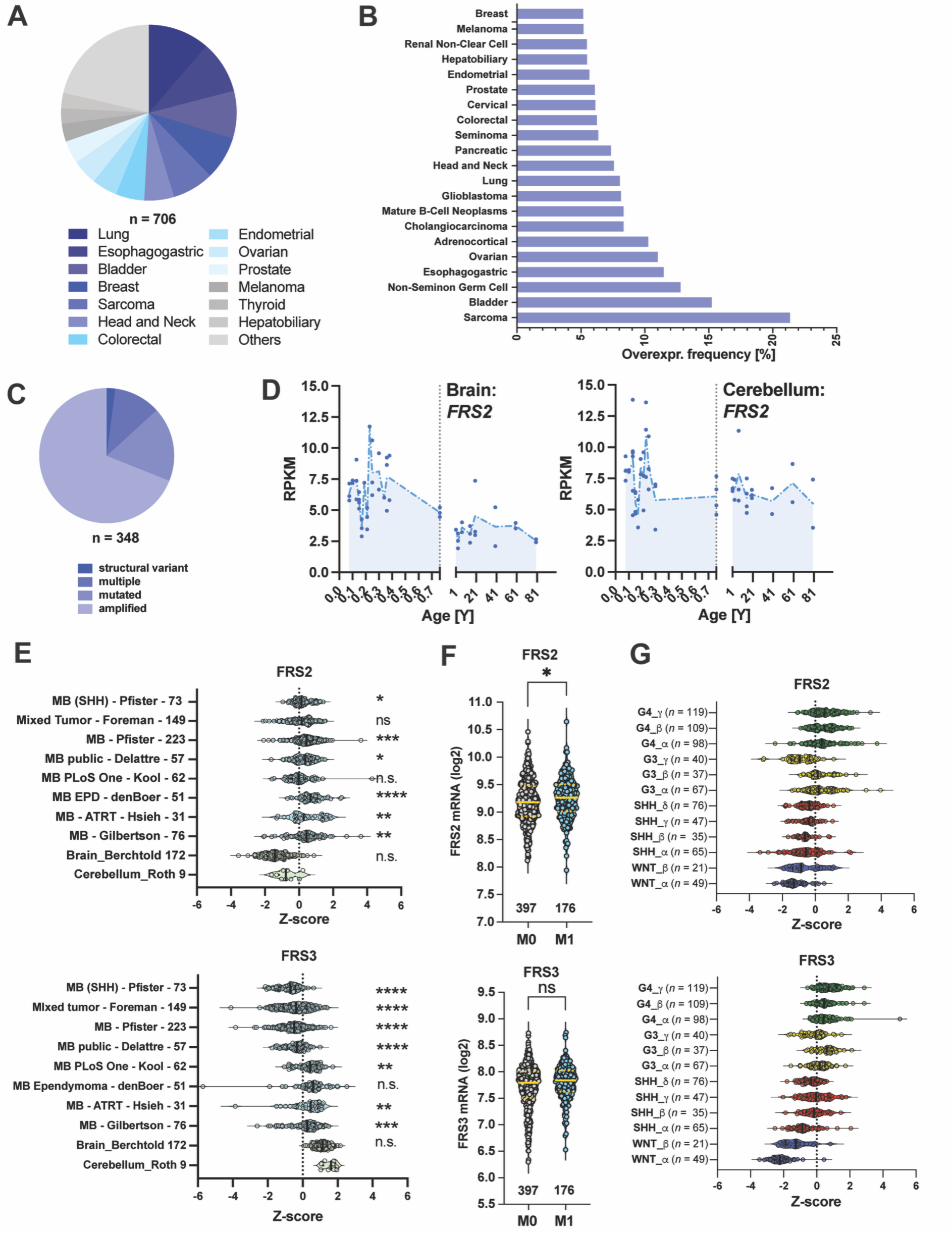
*FRS2* expression is elevated across multiple cancer types, including MB. **(A)** Pie chart showing the absolute number of TCGA PanCancer Atlas tumor samples with increased FRS2 levels. A z-score threshold of ±2, relative to mean expression in diploid samples, was applied as the cutoff for panels A– C. **(B)** Relative frequency of FRS2 overexpression across tumor diagnoses. **(C)** Absolute frequency of FRS2 genomic alteration types in cancer patients. **(D)** FRS2 expression from bulk RNA-seq data across the human lifespan in the brain (left) and cerebellum (right). The x-axis denotes age in years; the vertical dashed line marks birth. The y-axis shows expression values in reads per kilobase of transcript per million mapped reads (RPKM). **(E)** Comparative analysis of FRS2 and FRS3 mRNA abundance in normal cerebellum versus MB tumor tissue and other brain regions, performed on the Affymetrix U133P2 platform. Statistical comparisons were made using one-way ANOVA with Kruskal–Wallis and Dunn’s multiple comparisons tests; significance is indicated as follows: ns p ≥ 0.05, *p ≤ 0.0332, **p ≤ 0.0021, ***p ≤ 0.0002, ****p ≤ 0.0001. **(F)** FRS2 and FRS3 mRNA abundance in non-metastatic MB patients (M0) versus patients with tumor cells detectable in the cerebrospinal fluid (M1), assessed by unpaired two-tailed Mann–Whitney test (ns: p > 0.05). **(G)** FRS2 and FRS3 mRNA expression across the 12 MB molecular subtypes defined in the Cavalli dataset [21].

To assess differential expression in human lifespan, we profiled *FRS2* mRNA levels across time (prenatal to high age) in brain and cerebellum, using the Evo-devo atlas of bulk RNA-sequencing [36]. *FRS2* expression is highest during early development, both in brain and cerebellum and reaches a stable plateau after birth (Figure 1D). In MB patients, *FRS2* expression levels were increased in six out of eight investigated data sets compared to healthy cerebellum tissue (p < 0.05, Figure 1E), indicating a possible association of increased *FRS2* levels with MB. In contrast, *FRS3* was downregulated in MB tumors compared to healthy cerebellum and brain (p < 0.01, Figure 1E). A similar trend of increased *FRS2* in MB samples was observed in a batch normalized dataset comparing primary MB samples with normal cerebellum samples from a total of 23 databases [20] (Figure S1F). The analysis of the Cavalli dataset [21] consisting of 763 primary tumor samples further revealed increased *FRS2* expression in metastatic tumors M1 (tumor cells in cerebrospinal fluid) compared to M0 non-metastatic tumors (p < 0.05, Figure 1F), which was further corroborated by analyzing the batch normalized dataset (Figure S1G). Across MB subgroups, both *FRS2* and *FRS3* levels are higher in subgroups 3 and 4 (Figure 1G).

Although definitive evidence linking elevated FRS2 to cancer development or progression is still lacking, its upregulation in numerous malignancies warrants deeper investigation into its possible oncogenic roles.

### FRS2-driven motile phenotype is associated with FRS2 localization to leading-edge and cell-cell contacts

We previously demonstrated the relevance of FRS2 in FGFR-driven invasiveness in cell-based models of MB *in vitro* and in the tissue context [9,37]. The MB cell lines UW228 (SHH), ONS-76 and D425 (Grp3) display comparable FRS2 protein expression levels, while higher *FRS2* mRNA levels were observed in D425. (Figure 2A). We confirm high bFGF expression levels in the UW228 line only, which is consistent with previous data on *bFGF* mRNA expression across several MB cell-based models [9] (Figure 2A). Silencing FRS2 in DAOY and UW228 prevents bFGF-induced invasion [9], and elevating FRS2 expression increased bFGF-induced invasion in DAOY cells and reduce the sensitivity to a small molecule ligand of FRS2 [35]. This suggested that the augmented abundance of the FRS2 scaffold protein could be associated with cellular capabilities relevant for tumor growth and progression.

**Figure 2.**
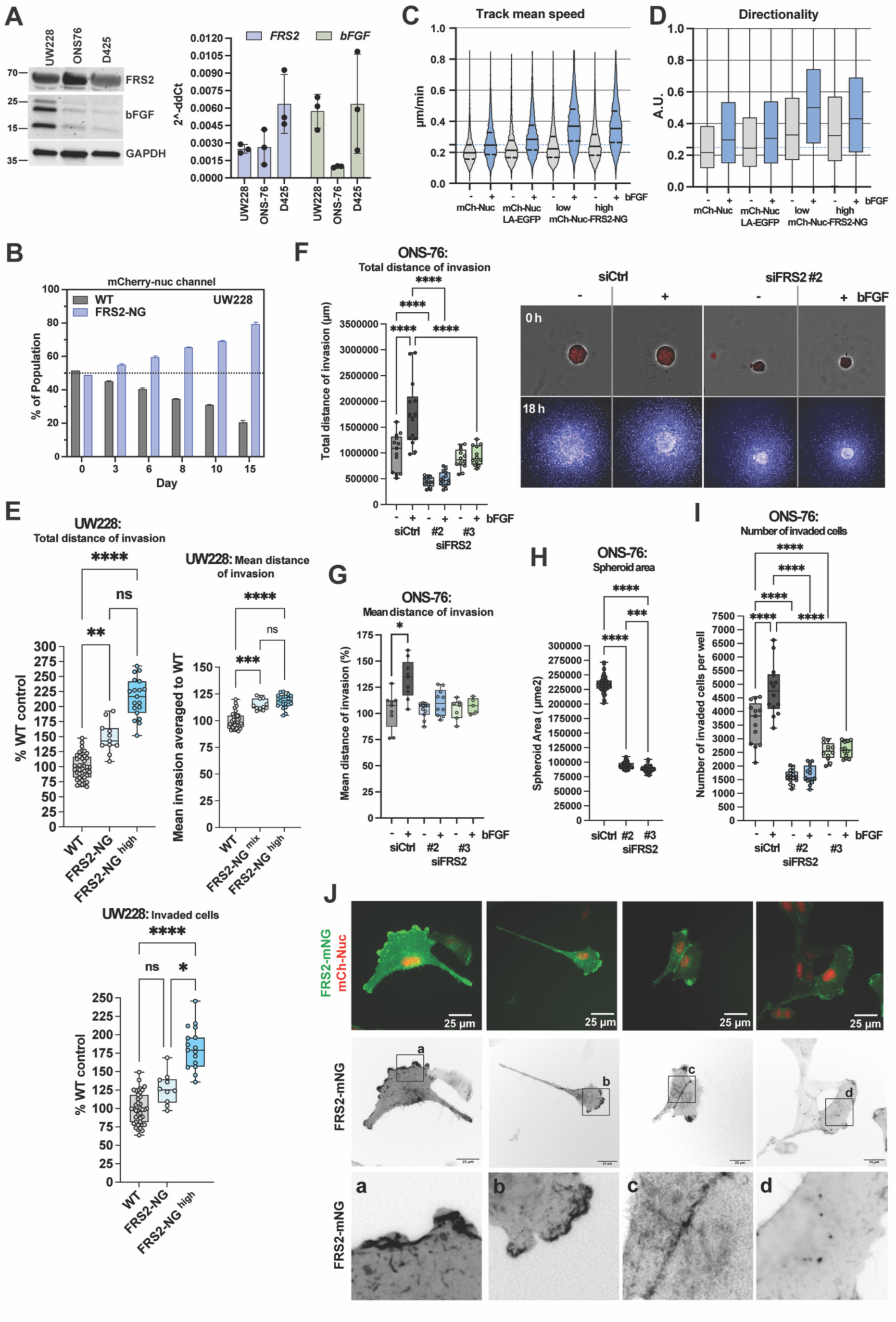
Elevated FRS2 expression is associated with increased proliferation, 2D migration, and 3D collagen invasion in MB cell lines. **(A)** IB and RT-qPCR analyses of FRS2 and bFGF expression in UW228, ONS-76, and D425 MB cells. **(B)** Proliferation competition assay in which wild-type UW228 and UW228 FRS2-mNG/mCherry-Nuc (mCh-Nuc) cells were grown in co-culture. The relative proportion of FRS2-mNG-positive cells was monitored over two weeks by flow cytometry using the mCh-Nuc signal. **(C)** Mean track speed and **(D)** directionality from live-cell tracking of UW228 mCh-Nuc (−bFGF: n = 4’601; +bFGF: n = 4’746), UW228 mCh-Nuc/LA-EGFP (−bFGF: n = 4’403; +bFGF: n = 4’729), and UW228 mCh-Nuc/FRS2-mNG cells (mixed −bFGF: n = 9’660; mixed +bFGF: n = 12’052; high FRS2 −bFGF: n = 8’195; high FRS2 +bFGF: n = 8’283), in the presence or absence of 50 ng/mL bFGF. **(E)** Invasiveness of wild-type UW228 versus FRS2-mNG-expressing UW228 cells assessed in serum-free medium over 24 h. Total invasion distance is calculated as the sum of individual invasion distances multiplied by the number of invading cells. Dots represent individual spheroids (each comprising >2,500 cells) from technical replicates of n = 1 experiment. **(F)** Left: Total invasion distance of ONS-76 cells transfected with siCtrl or siFRS2. Right: Representative images of spheroids prior to collagen I embedding (upper panels, brightfield with segmentation overlay) and 18 h after embedding (lower panels). Nuclei are labeled with Hoechst and computationally segmented. **(G)** Mean invasion distance of ONS-76 cells transfected with siCtrl or two independent FRS2 siRNAs. **(H)** Spheroid area (µm²) at the time of collagen matrix embedding, 48 h post-transfection. **(I)** Total number of invading ONS-76 cells after siRNA transfection. **(J)** Representative fluorescence live-imaging stills of UW228 cells co-expressing FRS2-mNG and mCh-Nuc on collagen I-coated surfaces. Panels a–d show magnified views of regions indicated by black rectangles in inverted greyscale images of the mNG channel, illustrating FRS2-mNG localization at the leading edge (columns 1–2), cell–cell contacts (column 3), and intracellular vesicles (column 4). Scale bar, 25 µm. Statistical comparisons were performed using one-way ANOVA with multiple comparisons correction; significance is indicated as follows: *p ≤ 0.05, **p ≤ 0.01, ***p ≤ 0.001, ****p ≤ 0.0001.

To explore this further, we first investigated whether FRS2 protein levels influence cell motility and invasion. As altered subcellular distribution dynamics might highlight relevant localization function relationships, we generated an UW228 MB cell line stably expressing FRS2 with a C-terminal mNeonGreen-tag (F-NG), to monitor FRS2’s subcellular distribution dynamics. The nuclear marker mCherry-Nuc was co-expressed to facilitate cell tracking. In addition, a population with uniformly high FRS2-mNG expression (F-NG^high^), to mimic FRS2 overexpression, was obtained by fluorescent activated cell sorting (FACS). The overexpression of FRS2-mNG in UW228 cells resulted in higher levels of Y436 phosphorylated FRS2 but did not correspondingly increase MAPK signaling, as assessed by IB detection of phosphorylation of ERK1/2 and AKT (Figure S2A). However, FRS2-mNG overexpression caused an estimated 9% increase in proliferation rate, resulting in the outgrowth of FRS2-mNG-m-Cherry-Nuc-positive (F-NG^high^) over wildtype (WT) UW228 cells (Figure 2B, S2B). UW228 F-NG and F-NG^high^ cells exhibited a significantly increased single-cell motility (p < 0.05, Figure 2C, S2C) and directionality (p < 0.005, Figure 2D, S2D) on collagen I coated plates compared to control UW228 cells transduced with LA-EGFP/mCherry-Nuc or mCherry-Nuc only. Increasing FRS2 expression increased total distance of invasion—quantified as the sum of individual cell invasion distances— and mean distance of invasion, which is a direct measurand of the capability of cells to migrate in collagen I gels (Figure 2E). This FRS2-mediated increase of invasiveness in UW228 cells was not dependent on exogenously added bFGF as no further increase in invasiveness was observed in the presence of exogenous bFGF (Figure S2E). We next examined the functional consequences of reduced FRS2 expression in the highly invasive ONS-76 cell line using RNA interference. ONS-76 cells readily invade collagen I gels, and bFGF stimulation further increases invasiveness by ∼30% (Figure 2F). FRS2 depletion (Figure S2F) caused a significantly decreased invasiveness, both in the absence and presence of bFGF (Figure 2F). Reduced invasiveness was associated with a decrease in average invasion speed (Figure 2G). The depletion of FRS2 caused a significant reduction of spheroid size (Figure 2F,H), which was associated with a reduced number of invading cells (Figure 2I). Together, these findings demonstrate the implication of FRS2 in the control of the invasive behavior of MB tumor cells, likely by influencing both migratory and proliferative capabilities of the cells.

Next, we determined the subcellular localization and distribution dynamics of overexpressed FRS2 by monitoring FRS2-mNG in UW228 cells. The cells were seeded in collagen IV-coated wells and maintained under regular growth conditions (see videos in Supplementary files 2-6). We observed the accumulation of FRS2-mNG at three distinct locations: the leading edge of migrating cells (Figure 2J, a, b), cell-cell contacts (Figure 2J, c), and intracellular vesicles (Figure 2J, d). Immunofluorescent analysis detecting endogenous FRS2 protein, which is enriched near the leading-edge membrane in UW228 wt cells (Figure S2G), confirmed the physiological distribution of FRS2-mNG.

Taken together, these results show that elevated FRS2 levels are associated with a motile and invasive phenotype in MB cells and enhanced proliferation, while no overactivation of canonical FGFR signaling was observed. Our data thus suggest that FRS2 may exhibit oncogenic activities, which are further increased by bFGF. The distinct localization patterns of FRS2 point towards its involvement in processes that regulate cell motility, adhesion, and intracellular transport.

### FGFR-induced recruitment of actin cytoskeleton regulators and translation initiators to FRS2

Since bFGF-expressing cells populate the tumor microenvironment (TME) in MB [9], we hypothesized that the local growth factor gradient could be involved in MB tumor control via FRS2-dependent FGFR signaling. *In vivo* and in cerebellar tissue slices, the repression of FGFR by the FGFR inhibitor BGJ389 reduces invasion [9]. The characterization of FGFR-induced phosphosites and of the FRS2 interactome could thus contribute to the identification of molecular mediators or pro-invasive FGFR-FRS2 signaling and of FRS2 function. Therefore, we used mass-spectrometry analysis to determine bFGF-sensitive protein phospho-sites and the FRS2 interactome.

Kinase-targeted phosphoproteomics analysis in bFGF-stimulated MCF-7 breast cancer cells revealed MAPK activation within 5 min of bFGF exposure, with a peak at 15 min [40]. 5 – 15 min stimulation with bFGF was found sufficient to induce FGFR signaling in MB cells [9,37]. We therefore first determined the minimal bFGF concentration necessary for maximal induction of FRS2 phosphorylation within 5 min, as we were interested in immediate, direct effects of FGFR signaling. We used the group 3 D425 MB cell model, which showed low baseline levels of FRS2, ERK, and AKT phosphorylation and maximal FRS2 phosphorylation within 5 min, 50 ng/mL (Figure S3A). We confirmed that FRS2 phosphorylation under these conditions was associated with the induction of ERK and AKT phosphorylation, indicating that intracellular FGFR signaling was activated (Figure S3A). Importantly, we also confirmed the bFGF-induced phosphorylation of the FRS2-interactors SHP2 and GAB1 (Figure 3A).

**Figure 3.**
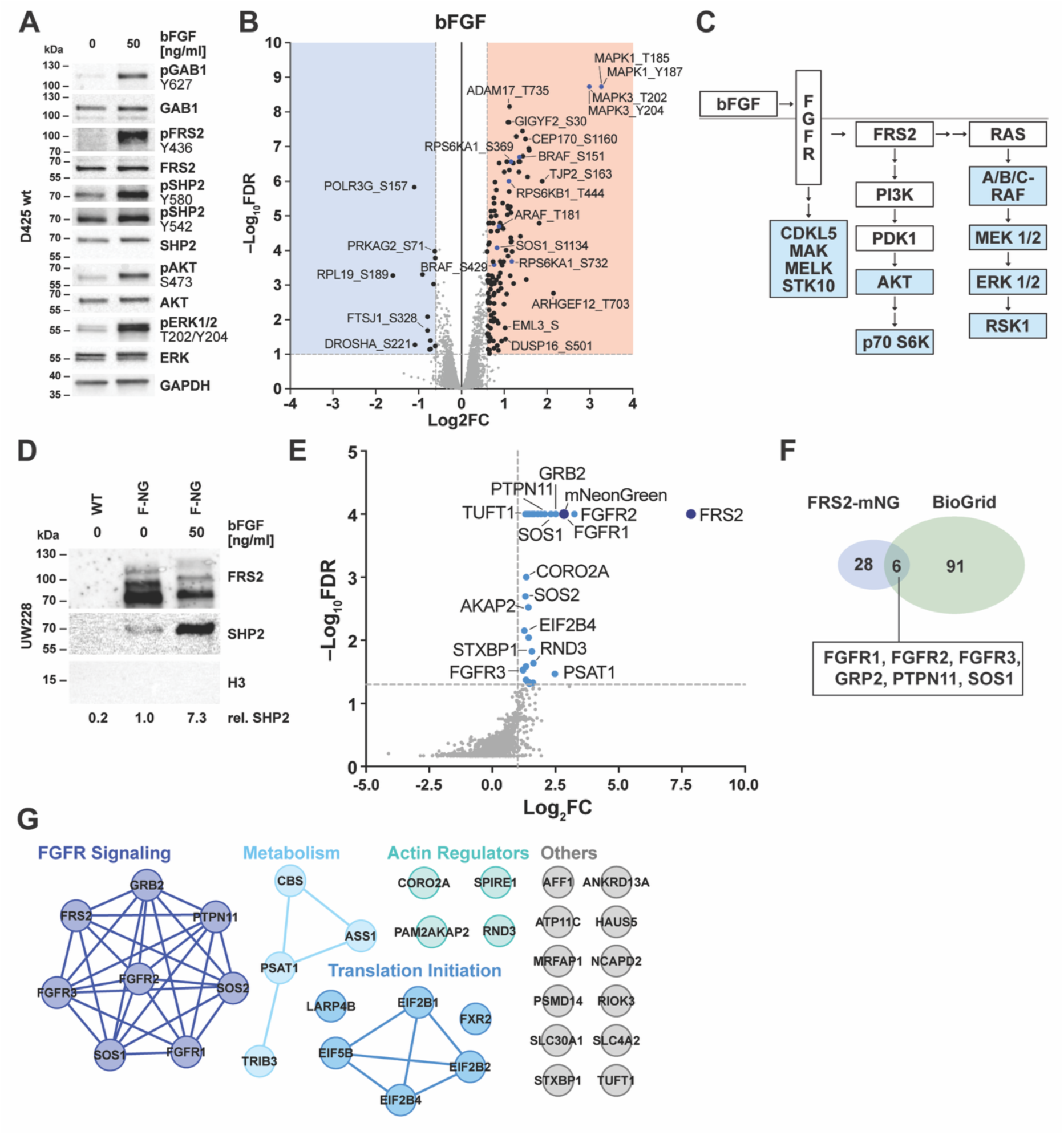
Canonical FGFR signaling is active in growth factor-stimulated MB cells and associated with the assembly of an FRS2-dependent signalosome. (**A**) IB analysis of D425 MB cells stimulated with 50 ng/mL bFGF for 5 min. The phosphorylation of FRS2 (Y_436_) and its interaction partners GAB1 (Y_627_) and SHP2 (Y_580_, Y_542_) as well as the activation of ERK1/2 (T_202_, Y_204_) and AKT (S_473_) pathways was detected using phosphospecific antibodies. (**B**) Vulcano plot of phosphoproteomic changes in of D425 cells after 5 min stimulation with 50 ng/mL bFGF (*n*=3 biological replicates). Significantly downregulated sites (FDR ≤ 0.1, log_2_FC ≤ –0.6) are highlighted in blue and significantly upregulated (FDR ≤ 0.1, log_2_FC ≥ 0.6) in red. Phosphosites belonging to FGFR pathway members are indicated in blue. (**C**) Schematic of FRS2-dependent canonical FGFR signaling, encompassing the MAPK/ERK and PI3K/AKT branches. Kinases with bFGF-sensitive phospho sites are shown in blue. (**D**) IB of the eluate fraction of mNeonGreen-Trap IP performed in wild-type UW228 and FRS2-mNG-expressing UW228 cells (± 50 ng/mL bFGF, 5 min). (**E**) Vulcano plot of FRS2-mNG co-precipitated proteins identified by mass spectrometry. Significantly enriched proteins (FDR ≤ 0.05, log2FC ≥ 1) are highlighted in blue (*n*=3 biological replicates). **(F)** Venn diagram comparing proteins enriched in the FRS2-mNG co-IP (excluding mNeonGreen but including FRS2) with previously reported FRS2 interactors catalogued in the BioGRID database. **(G)** STRING network analysis of FRS2-mNG co-precipitated proteins, highlighting distinct interaction clusters among predicted FRS2-associated candidates.

To assess global alterations induced by FGFR signaling, we performed a phospho-proteomic analysis under identical conditions and identified 143 bFGF-regulated phosphorylation sites on 112 unique proteins (FDR ≤ 0.1, abs(log_2_FC) ≥ 0.6) including 10 kinases (Figure 3B, Supplementary file 7). To check for the sensitivity of our cutoff, we also applied a more stringent FDR of ≤ 0.05, which reduced the number of detected proteins with altered phosphorylation by only 10 (five down: DROSHA, TNF2, MAP1B, FASTKD2 and TESK, five up: by RECQL4, RAD51C, PTGFRN, TULP4 and TINF2). The key members of the canonical FGFR signaling pathway were among the phospho-proteins detected under bFGF-stimulated conditions (Figure 3C). Our analysis also uncovered the human kinases cyclin-dependent kinase-like 5 (CDKL5), male germ cell-associated kinase (MAK), maternal embryonic leucin zipper kinase (MELK) and serine/threonine kinase 10 (STK10) as novel putative bFGF-sensitive effectors, potentially implicated in FGFR-driven cellular processes. Hemoglobin subunit alpha (HBA1/2) and tropomodulin-2 (TMOD2) were the only proteins with slightly increased total protein expression levels (FDR ≤ 0.1, abs(log_2_FC) ≥ 0.6) (Figure S3B), arguing against a rapid change in protein abundance after short term bFGF stimulation.

To study the interactome of FRS2 in the FGFR-activated state, we performed immune precipitation (IP) of FRS2-mNG from bFGF-stimulated UW228-FRS2-mNG cells. We detected the binding of the known FRS2 interactor SHP2 to FRS2-mNG and confirmed its increased interaction in response to bFGF stimulation (Figure 3D). Anti-mNG affinity purification followed by mass spectroscopy identified 37 significantly enriched putative interactors of FRS2 in UW228-FRS2-mNG cells (FDR ≤ 0.05, log2FC ≥ 1) compared to anti-mNG affinity purification in control (UW228 wt) cells (Figure 3E, Table 1, Supplementary file 8). Importantly, the co-precipitation of known FRS2 interactors including FGFR1-3, GRB2, SHP2 or SOS1/2 indicate the correct integration of the FRS2-mNG fusion protein into the FGFR signaling pathway. Six of the 34 co-precipitated proteins have been previously documented as FRS2 interactors in the BioGrid data base (Figure 3F), whereas 28 proteins can be considered as novel candidate interactors of FRS2. String-network analysis of the co-precipitated proteins predicts an FGFR signaling cluster, and three additional clusters associated with metabolism, actin cytoskeleton regulation and translation initiation (Figure 3G).

bFGF stimulation thus activates canonical FGFR signaling in MB cells, induces the phosphorylation, and our data furthermore predict the recruitment of proteins involved in actin cytoskeleton regulation, translation initiation and metabolism in bFGF-stimulated cells.

### Regulators of the actin cytoskeleton and cell-cell contacts constitute the proximal proteome of FRS2

To analyze the molecular environment of FRS2 in an intact cellular context, we established an engineered ascorbate peroxidase 2-based (APEX2) proximity labeling protocol for the FRS2 protein in UW228 cells. We used the UW228 SHH MB model as we determined FRS2-dependent motile capabilities (Figure 2C-E), established FRS2 localization dynamics (Figure 2J), and determined the FRS2 interactome (Figure 3G) in this cell line. APEX2 allows for rapid intracellular biotinylation of proteins in a range of 10 – 20 nm of the fused biotin ligase [22,38] (Figure 4A). With this approach, we aimed at the identification of proteins constitutively associated with FRS2 and of proteins that could associate with FRS2 in a bFGF stimulation-dependent manner.

**Figure 4.**
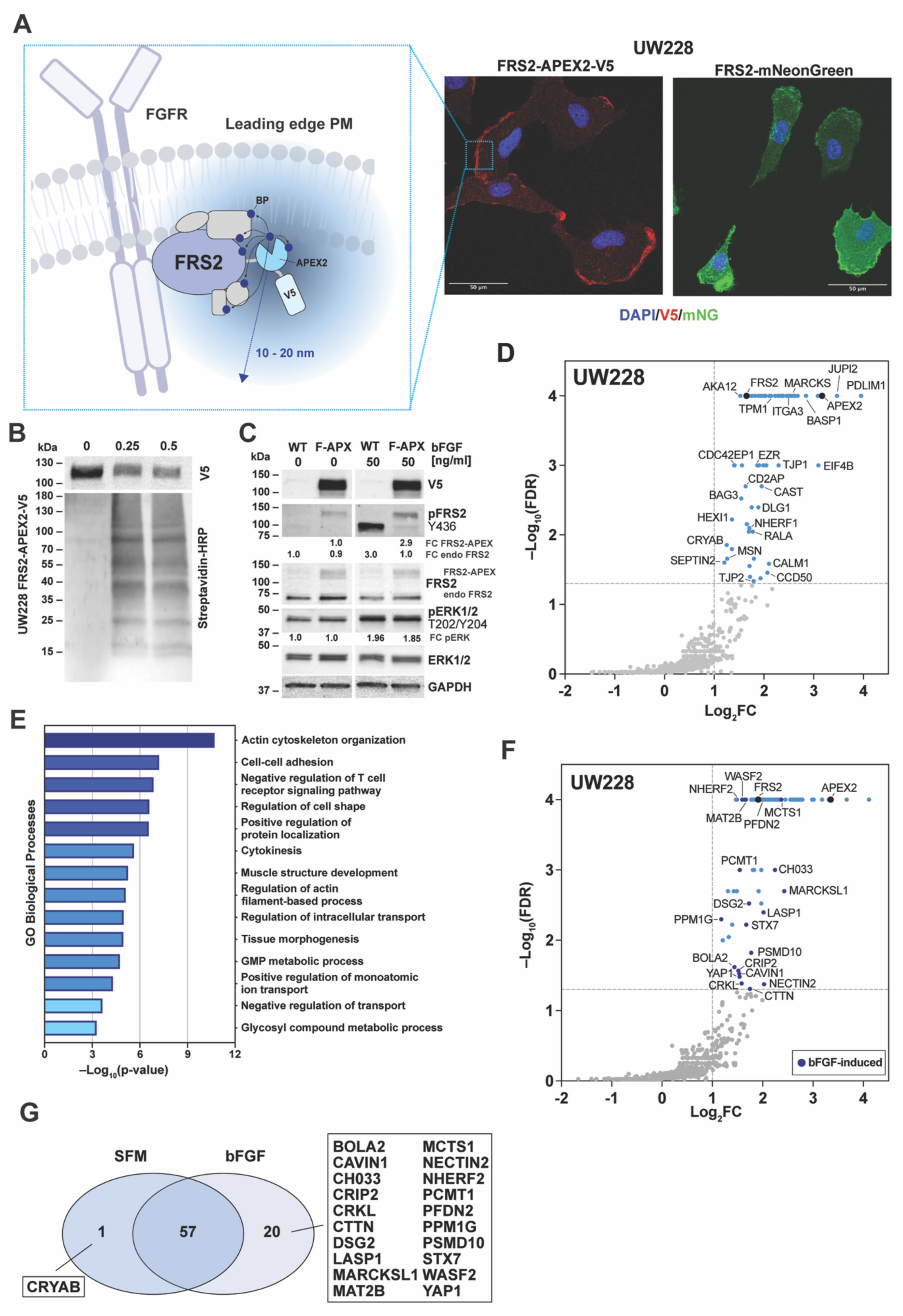
Identification of proteins in FRS2 proximal proteome using APEX2-based proximity labeling. (**A**) Left: Cartoon depicting the fusion of an APEX2 peroxidase enzyme to the C-terminal end of FRS2 and the assumed proximity labeling range using biotin-phenoxyl radicals (BP). Right: Immunofluorescence analysis of subcellular localization of FRS2-APEX2-V5 and FRS2-mNG in UW228 cells. (**B**) Comparative IB analysis of FRS2-APEX2-V5 expression and Biotin-HRP detection of biotinylated proteins after H_2_O_2_-treatment. (**C**) IB analysis of bFGF-induced phosphorylation of FRS2 (Y_436_) and ERK1/2 (T_202_, Y_204_) in wt and FRS2-APEX2-V5 (F-APX)-expressing UW228 cells. Fold change (FC) of endogenous FRS2 and F-APX phosphorylation and of pERK are shown. WT and F-APX without bFGF stimulation were used as baseline to calculate FC after bFGF stimulation. (**D**) Vulcano plot of proteins enriched in FRS2-APEX2-based proximity labeling assay in serum-free culture conditions (SFM). Significantly enriched proteins (FDR ≤ 0.05, log_2_FC ≥ 1) are shown in light blue. The bait control proteins FRS2 and APEX2 are shown in black (*n*=3 biological replicates). (**E**) Over representation analysis depicting enriched gene ontology terms (biological processes) of FRS2-APEX2-V5 labeled proteins. (**F**) Vulcano plot of FRS2-APEX2-V5 labeled proteins in bFGF-stimulated conditions (50 ng/mL bFGF stimulation for 5 min). Significantly enriched proteins (FDR ≤ 0.05, log_2_FC ≥ 1) are shown in blue (*n*=3 biological replicates). Dots of proteins significantly enriched in the presence of bFGF are colored dark blue. The bait control proteins FRS2 and APEX2 are shown in black. (**G**) Venn diagram of proximity-labeled proteins detected in serum-free (SFM) and bFGF-stimulated (bFGF) conditions. FRS2 and APEX2 are included (overlapping fraction).

We stably expressed FRS2-APEX2-V5, which localizes, comparably to FRS2-mNG, to the leading edge of migrating cells (Figure 4A, S4A). We next confirmed that H_2_O_2_-induced protein biotinylation in UW228 FRS2-APEX2-V5 cells (Figure 4B), and that the FRS2-APEX construct was expressed at a level comparable to the endogenous FRS2 protein (Figure 4C). Importantly, bFGF stimulation induced phosphorylation of Y436 in FRS2-APEX2-V5 and downstream ERK1/2 activation comparable to the WT control (Figure 4C, S4B). Furthermore, a biotinylated peptide encompassing the FGFR juxtamembrane region affinity-precipitated the FRS2-APEX2-V5 protein, confirming its ability to bind native FGFR receptors (Figure S4C).

We found direct processing of pulled-down APEX2-labeled proteins without freezing (Figure S4D) and lysis buffer exchange via acetone precipitation to remove excess biotin-phenol substrate (Figure S4E) essential to ensure optimal pull-down efficiency. As the experimental conditions, we opted for rapid proximity labeling (3 min) and comparing serum-starved and bFGF-stimulated cells, to capture both constitutive and FGFR activation-dependent changes in the proximal proteome of FRS2. As a negative control, we used UW228 cells expressing FRS2-mNeonGreen.

Under serum starvation and in the absence of bFGF, we detected a total of 58 significantly enriched unique proteins (FDR < 0.05, log2FC ≥ 1) in the proximity of FRS2 (Figure 4D, Table 2, Supplementary file 9). We also detected the biotinylation of FRS2 and APEX2. Over Representation Analysis (ORA) predicted significant enrichment for proteins involved in F-actin dynamics, the regulation of cell shape, cell-cell adhesion, and protein localization among others (Figure 4E). Identified candidates of the FRS2 proximal proteome included plasma membrane-associated proteins (e.g. BASP1, MARCKS, EZR, MSN), actin filament regulators (e.g. PDLIM1, TPM1/2, CALD1), cell junction proteins (TJP1/2) and vesicle-associated proteins (GNPDA1, DLG1). The described subcellular distribution and functions of these interactors are consistent with the observed localization of FRS2-mNG (Figure 2J). Following bFGF stimulation, we identified an additional 20 candidate proteins recruited to the proximity of FRS2 in a stimulation-dependent manner (Figure 4F,G, Table 3, Supplementary file 9). These included multiple plasma membrane-associated proteins (e.g. WASF2, MAT2B, NHERF2), proteins in cell junctions (DSG2, NECTIN2), and actin filament-binders (MARCKSL1) (Figure S4F). Only three of all identified candidates (EZR, MARCKS, STX7) have been documented as potential FRS2 interactors in the BioGrid data base (Figure S4G).

In summary, we confirmed known interactors and describe 75 novel candidate proteins potentially associated with FRS2 function. For 20 of these proteins, the association is bFGF-dependent. Our data suggest the association of FRS2 with proteins involved in the regulation of the actin cytoskeleton, cell junctions, and intracellular transport in MB cells.

### Interaction proteomics predict FRS2 coupling to cytoskeleton organization, cell junction assembly and intracellular transport

The phosphoproteome- and APEX2-predicted proteins could represent the molecular environment of FRS2 that may weakly, transiently, or indirectly interact with FRS2. Notable in this group are TJP1/2 and EIF4B, which are predicted components of the proximal proteome of FRS2 and harbor bFGF-regulated phospho-sites (Figure 3B, 4D).

String-network analysis of the predicted FRS2-interacting or proximal proteins revealed a complex network in the vicinity of FRS2 (Figure 5A), with the cluster of canonical FGFR signaling proteins associated with a network of cytoskeleton regulators and cell junction proteins. Distinct groups of proteins involved in translation initiation and intracellular transport-associated proteins were additionally predicted.

**Figure 5.**
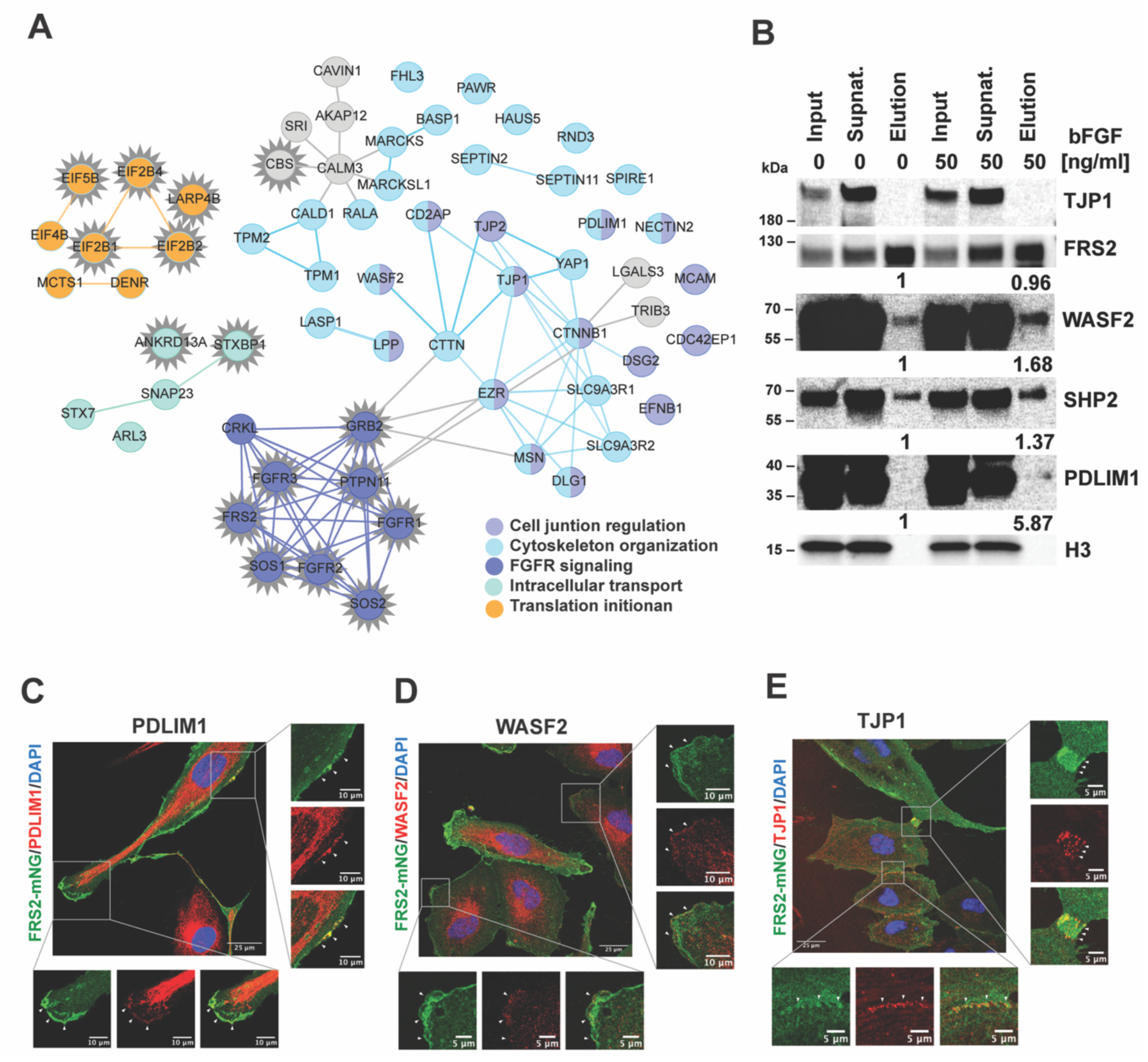
FRS2 connects FGFR signaling to cytoskeletal organization, intracellular transport, and translation initiation. **(A)** STRING protein network analysis of FRS2 interactor candidates identified by co-immunoprecipitation (star-shaped nodes) and APEX2 proximity labeling. **(B)** IB analysis of proteins co-immunoprecipitated with FRS2-mNG. Input, post-IP supernatant, and eluate fractions were probed with the indicated antibodies. Numbers below each panel represent the fold change in integrated density of the eluate from bFGF-stimulated cells relative to unstimulated controls. (**C–E)** Immunofluorescence analysis of PDLIM1 (C), WASF2 (D), and TJP1 (E) in UW228 cells expressing FRS2-mNG.

To validate a small subset of predicted proximal proteome components of FRS2, we confirmed FRS2-APEX2-mediated biotinylation of WASF2, TJP1 and YAP1 by streptavidin pull-down and IB (Figure S5A). EZR and PDLIM1 could not be detected using this approach. As an alternative, we performed FRS2-mNG pull-down assays and confirmed FRS2 interaction with WASF2 and with the positive control SHP2 (Figure 5B). Notably, bFGF stimulation enhanced the predicted interaction with both WASF2 and PDLIM1 and with the positive control SHP2. To score relevant interactors, we compared the predicted interactors from the phosphoproteome analysis, the APEX2-labeling, and the FRS2-mNG pull-down with the previously compiled FRS2-interactors documented in the BioGrid database (Figure S5B). The FRS2-mNG pull-down predominately captured interactors of the canonical FGFR signaling pathway (FGFR1-3, GRB2, PTPN11, SOS1). These well-characterized interactors are listed in the BioGrid database, confirming the functionality of the FRS2-mNG construct in our cellular models. The APEX2-detected proteins overlap more with the bFGF-induced phospho-proteome data set, and include MARCKS and EZR, both of which are documented in the BioGrid database.

We next analyzed the subcellular localization of PDLIM1, WASF2, and TJP1 by immunofluorescence analysis in UW228 FRS2-mNG cells. We observed all three proteins in proximity of FRS2-mNeonGreen at the cell periphery, specifically within lamellipodia (PDLIM1, WASF2) and at cell–cell junctions (TJP1) (Figure 5C–E), suggesting that their interaction with FRS2 may occur in these compartments. Mapping the biotinylated peptides identified by the APEX2 proximity labeling onto the AlphaFold structures of WASF2, PDLIM1, and TJP1 (Figure S5C-E) [39] revealed an enrichment of biotinylation in regions predicted as intrinsically unstructured or as short α-helices. For TJP1, multiple biotinylated peptides were located in close proximity to its predicted PDZ and SH3 domains, regions critically implicated in phase transition, subcellular localization and accumulation of TJP1 [40].

These experiments thus further substantiate the potential interaction and co-localization of WASF2, PDLIM1 and TJP1 with FRS2 or a FRS2-associated protein complex in cells and point towards FRS2-mediated processes in cytoskeletal regulation and cell junction control. However, no clear functional link between FRS2-mediated motile processes and these cytoskeleton-associated proteins at single cell level in MB cells could be established.

### FRS2 enables bFGF-induced TJP1 clustering

Clustering of the predicted FRS2 interactor TJP1 in membrane associated protein complexes is enabled by distinct phosphorylation events in TJP1 [40]. To investigate the relevance of FRS2 in this context more broadly and independent of the MB tumor cell setting, we assessed TJP1 localization in wild-type (HAP1 wt) and FRS2 knockout (HAP1 FRS2 KO) HAP1 cells. In HAP1 FRS2 KO cells, *FRS2* is not detectable at mRNA level, confirming the absence of FRS2 in these cells (Figure S6A). *FRS2* knockdown did not alter the expression levels of *FRS3* and *FGFR1*, *3* and *4*, but is associated with an upregulation of *FGFR2*. In wt cells, *FRS2* transcript levels were significantly higher than the *FRS3* levels (Figure S6B).

Phenotypically, HAP1 FRS2-KO cells displayed reduced cell–cell clustering, with individual cells remaining more distinctly separated compared to the compact clusters observed in HAP1 wt cells (Figure 6A, overlay). To test whether altered TJP localization in HAP1 FRS2 KO cells could be associated with this phenotype, we examined the bFGF-dependent localization of TJP1 by immunofluorescence (Figure 6A). In HAP1 wt cells, bFGF-induced, distinct TJP1 aggregation at basal sites of the cells was observed (Figure 6A,B). This phenotype was reduced in HAP1 FRS2 KO cells, where TJP1 failed to form large aggregates (Figures 6A-C, S6C). IB analysis revealed no differences in TJP1, WASF2 or PDLIM1 expression in HAP1 wt and HAP1 FRS2 KO cells, excluding reduced protein abundance as possible cause of reduced clustering of TJP1 (Figure S6D).

**Figure 6.**
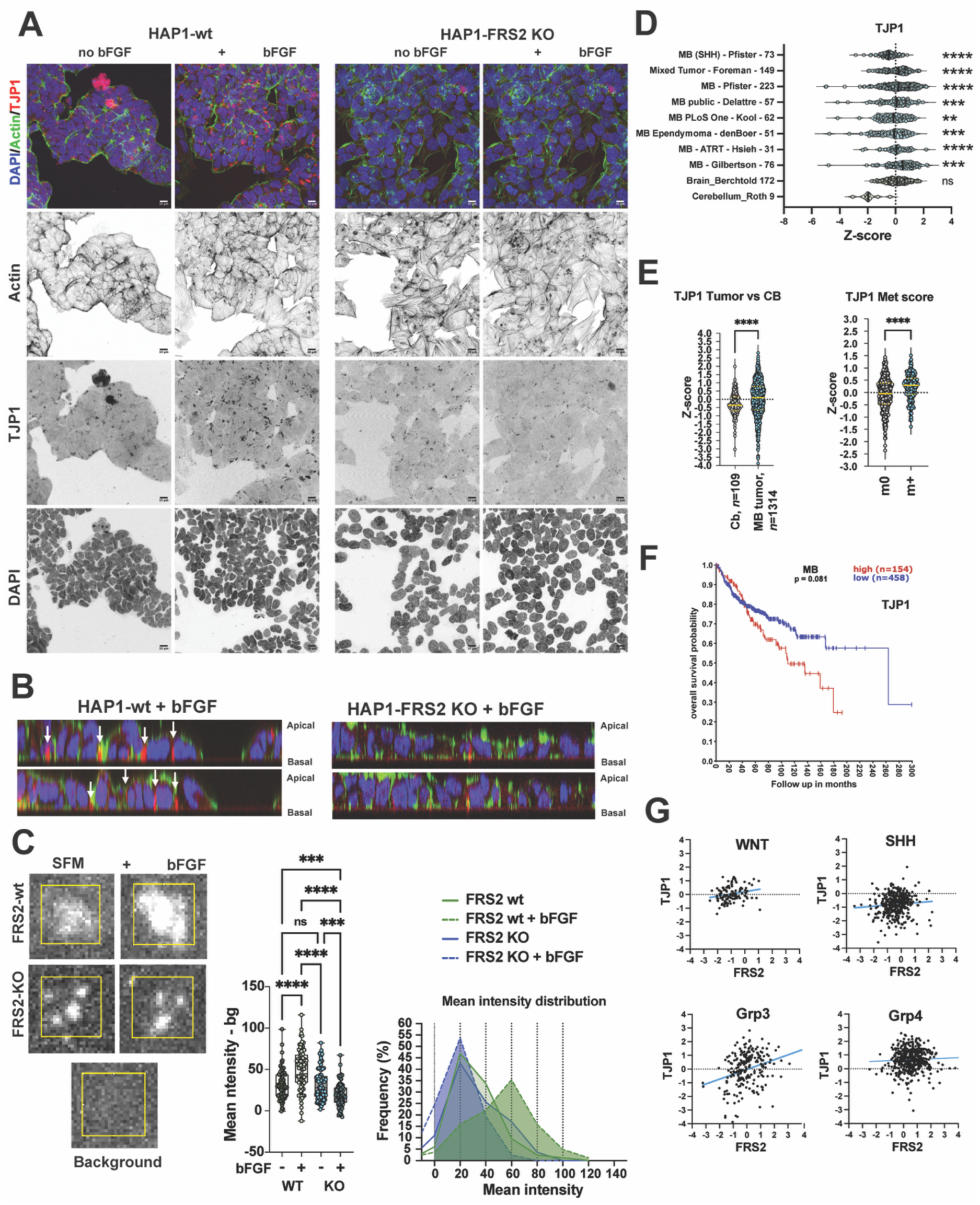
FRS2 is required for TJP1 clustering following bFGF stimulation. **(A)** Confocal immunofluorescence imaging of HAP1 FRS2-wt and HAP1 FRS2-KO cells stimulated with 50 ng/mL bFGF for 1 h. Green: F-actin (phalloidin); red: TJP1; blue: DNA (DAPI). Inverted greyscale images of each fluorescence channel are shown separately. **(B)** Orthogonal cross-section through confocal image stacks of HAP1-wt and HAP1 FRS2-KO cells stimulated with bFGF, as shown in (A). White arrows indicate basally clustered TJP1 between adjacent cell nuclei. **(C)** Quantification of TJP1 clustering. Left: The FRS2-APEX2-V5 was used as bait and included Clustered TJP1 signal intensity was quantified as the mean pixel intensity within a 20 × 20 pixel region of interest encompassing each TJP1 cluster (n = 81 per condition; n = 1 experiment). Center: Box plots showing the median and distribution of measurements across the four conditions. Statistical significance was assessed by one-way ANOVA with Kruskal–Wallis test. Significance is indicated as follows: ns p ≥ 0.05, ***p ≤ 0.0009, ****p <0.0001. Right: Frequency distribution of mean intensities for each condition. **(D)** Z-scores of *TJP1* mRNA expression in two control datasets (normal brain regions and cerebellum, shown in green) and six primary MB patient datasets (shown in blue), analyzed on the Affymetrix U133P2 platform. Significant differences relative to the healthy cerebellum control were assessed by one-way ANOVA with Kruskal–Wallis and Dunn’s multiple comparisons tests; significance is indicated as follows: ns p ≥ 0.05, *p ≤ 0.0332, **p ≤ 0.0021, ***p ≤ 0.0002, ****p ≤ 0.0001. **(E)** Left: Z-scores of *TJP1* expression in cerebellar control (n = 109) and MB tumor samples (n = 1’314). Right: Z-scores of *TJP1* expression in non-metastatic MB patients (M0) versus patients with tumor cells detectable in the cerebrospinal fluid (M+; Swartling dataset, GSE124814, n = 1’641). Statistical significance was assessed by unpaired two-tailed Mann–Whitney test: ns p ≥ 0.05, ****p ≤ 0.0001. **(F)** Kaplan–Meier curves of overall survival in MB patients stratified by high (red) or low (blue) *TJP1* tumor expression, derived from the PanCancer database. **(G)** Correlation plots of *FRS2* versus *TJP1* mRNA expression z-scores across MB subgroups, with linear trendlines shown in blue.

To assess the potential clinical relevance of TJP1 in MB, we explored its expression levels and correlations with clinical parameters in publicly available datasets. We found that *TIP1* and *PDLIM1* but not *WASF2* expression are significantly higher in MB tumors compared to normal cerebellum (Figure 6D, S7A,B). Moreover, tumors from patients with metastatic MB cells detected in the cerebrospinal fluid (M1) displayed significantly higher *TJP1* and lower *WASF2* expression compared to non-metastatic tumors (M0) (Figure 6E, S7C,D). A correlation between higher *TJP1* or *PDLIM* expression and poorer overall survival of MB patients was observed (Figure 6F, S7E,F). The expression of *FRS2* and *TJP1* correlate positively in more metastatic Grp3 MB tumors, whereas *FRS2* and *PDLIM1* correlate negatively in this subgroup (Figure 6G, S7G).

In summary, our data indicate that FRS2 enables distinct bFGF-induced subcellular localization of TJP1, underscoring its role as a versatile scaffold that links FGFR signaling to downstream effector proteins. Transcriptomic analyses of primary MB tumor samples further suggest that TJP1, WASF2, and PDLIM1 may contribute to MB biology, with increased *TJP1* and *PDLIM1* being associated with poor prognosis.

## Discussion

FRS2 is a scaffold protein well-characterized in its function to assemble the FGFR signalosome, thereby linking receptor activation to MAPK/ERK and PI3K/AKT signaling pathways [5,7,8,33,41]. Its structure consists of a N-terminal PTB domain that binds the juxtamebrane region of FGFRs and a C-terminal unstructured region with multiple docking sites for several known interactors such as GRB2 [6] and SHP2 [8]. Whereas its role in FGFR signaling has been extensively studied, the components of the FRS2 signaling complex and its ability to bridge receptor activation to additional cellular processes remained incompletely understood. By investigating the expression, subcellular localization and the interactome of FRS2 in MB tumor cells, our study revealed potential molecular effector candidates associated with its role in promoting aggressive cellular phenotypes. This is particularly relevant in the context of MB, where we found elevated levels of FRS2 to correlate with a more motile and invasive phenotype *in vitro* and metastasis in patients. Whereas FRS2 expression levels alone may contribute to these phenotypes, pathway activation by bFGF enables the phosphorylation of proteins involved in the regulation of cell-cell contacts, the actin cytoskeleton and the initiation of translation. Collectively, by identifying novel components of the FRS2 signalosome, our study provides insights into how FRS2 could fine-tune the biological outcome of FGFR signaling and expands the current knowledge of FRS2’s role in FGFR-driven cancers.

Approaches using co-IP have previously been used to extract stable and high-affinity interactions for characterizing the interactome of FRS2 [6–8,15,17,32,42–46]. These studies mostly identified members of the canonical FGFR signaling pathways due to the limitations of classical pull-down methodologies to predominately capture strong and direct FRS2 interaction partners. By leveraging APEX2-based proximity labeling and classical pull-down methodologies, we identified 112 (IP + APEX) candidate proteins that may either directly interact with FRS2 or are present in associated protein complexes. Whereas the resulting FRS2 signalosome in MB tumor cells includes interactors previously described in other cellular systems such as FGFRs, GAB1, SHP2 and SOS1/2 [6–8,32,33,47], our pull-down approach in FGFR activated state identified 27 proteins that have not been described as potential FRS2 interactors so far. Additionally, we identified 78 transient or weak FRS2 interactor candidates detected only by proximity labeling. The labeling range of the APEX2-based approach is limited by the diffusion radius of short-lived biotin-phenoxyl radicals, usually resulting in labeling ranges of 10-20 nm [38]. The resulting identification of transient or weak interactions in the proximal proteome of FRS2 provides a more comprehensive insight in signalosome dynamics. Furthermore, as the labeling occurs in living cells, interactions that are disrupted through the lysis step in the co-IP approaches are preserved. This could explain why the proximity labeling assay detected proteins involved in dynamic cellular processes such as cytoskeletal remodeling, membrane dynamics and translation regulation that have not been detected with conventional methodologies so far. Importantly, 20 of these proteins were preferentially recruited to the proximal proteome of FRS2 in response to bFGF-stimulation, substantiating the notion of activity-dependent reorganization of the FRS2 signalosome. The presence of proteins with no established roles in canonical FGFR signaling suggests that FGFR-induced structural reorganizations of the FRS2 signalosome may be involved in actin regulation, translation initiation and metabolism. Paradigmatic for this is the interaction of FRS2 with the actin regulator WASP family protein member 2 (WASF2/WAVE2), which hints towards FRS2 implication in cell motility through modulating leading-edge extension dynamics [48,49]. WASF2 activity depends on PtdIns(3,4,5)P_3_ [50] and its phosphorylation at the leading edge by growth factor activated ERK [48] to promote the F-actin nucleation activity of the ARP2/3 complex [49]. We previously detected bFGF-induced activation of ERK at or near the plasma membrane [9].However, as we did not detect WASF2 in our phospho-proteome dataset, we cannot conclude on the phosphorylation status of FRS2-associated WASF2 in response to bFGF stimulation. However, the predicted FRS2 association with WASF2 and PDLIM1 corroborates the role of FRS2 in actin reorganization and lamellipodia formation [51–56] and suggests the mechanistic implication of FRS2 in WASF2-ARP2/3 control of leading-edge F-actin reorganization. Our data also confirmed FRS2 interaction with the negative regulator of RHOA activity, Rho-related GTP-binding protein RhoE (RND3) [15]. While the functional consequences of this interaction remain to be elucidated, increased RHO activity was observed in bFGF-stimulated cells with reduced FRS2 expression [9], suggesting the presence of a regulatory feedback loop that may involve FRS2 coupling to RHO signaling through RND3.

bFGF-sensitive phosphorylation sites in proteins involved in cell adhesion and migration, including protein Shroom3 (SHROOM3), afadin (AFDN) and membrane-associated guanylate kinase WW and PDZ domain-containing protein 3 (MAGI3) further underscore the notion of FRS2 controlling F-actin dynamics and cell motility. Potentially druggable candidates in the bFGF-sensitive phosphoproteome are the kinases CDKL5, MAK, MELK, and STK10. No report on FGFR or FRS2 interaction with CDKL5, MELK, and STK10 was found in literature, whereas MAK has been identified as a direct interactor and signaling mediator of FGFR3 [57], suggesting that FGFR activation of MAK involves FRS2.

In the predicted interactome of FRS2, we detected proteins involved in tight junction, cell adhesion or actin regulation, which also harbor bFGF-sensitive phosphorylation sites. These include MARCKS, TJP2 and EIF4B. Although we could not confirm altered TJP1 phosphorylation after bFGF stimulation in MB tumor cells in our phosphoproteomic dataset, the phosphorylation of the FRS2 interactors TJP1, LPP, CTNNB1 CD2AP and DSG2 has already been reported in bFGF-stimulated human embryonic stem cells [4]. In that model, increased activation of SRC family kinase was suggested to maintain an undifferentiated embryonic stem cell state by promoting cytoskeletal and actin-depending processes. SRCs are recruited to active FGFRs in a FRS2-dependent manner [58], and the assembly of SRC substrates on the FRS2 signalosome might enable their indirect regulation by SRC family kinases recruited to activated FGFRs. Consistently, FRS2 is required for bFGF-induced clustering of TJP1 in HAP1 cells, indicating a potential contribution of FRS2 in regulating SRC kinase control of critical regulators of actin dynamics and cell-cell interaction. However no clear functional link at single cell level between FRS2-mediated motile processes, TJP1 localization and the cytoskeleton-associated proteins identified as FRS2 interactor candidates could be established in MB cells so far. While recent studies have shown that junctional proteins can modulate FGFR signaling and contribute to tumor progression [59], our findings extend this model by implicating the canonical FGFR scaffold protein FRS2 in the regulation of TJP1 distribution. Given the scaffold properties of both proteins and their localization to plasma membrane-proximal compartments, their interplay is likely to be context dependent. In physiological epithelia, FGFR-FRS2 activation may enable tissue remodeling and repair by modulating junctional stability. A similar mechanism could also promote epithelial to mesenchymal transition in epithelial tumors with overactivated FGFR signaling.

An additional protein cluster in the interactome of FRS2 is composed of proteins involved in the regulation of translation (EIF2B1-2, EIF2B4, EIF4B, EIF5B, MCTS1, DENR, LARP4). Consistent with bFGF control of translation is the bFGF-inducible phosphorylation of serine and threonine residues on EIF4B (S_230_), EIF2A (S_526_) and LARP1 (T_376_). This could indicate ligand-dependent coupling of FGFR signaling to translational control via FRS2 in MB tumor cells, possibly through the FGFR downstream kinases RSK, S6K1 and AKT [60–62].

Spatially controlled activation of FGFR promotes cell motility through induced lamellipodia formation [3]. LARP family proteins have been described in regulating mRNA localization in these structures, to increase local translation and sustain cell migration and invasion [52], independent of the direct control of F-actin modulating proteins. Our proteomic and microscopy data indicate that FGFR-FRS2-enriched compartments in lamellipodia could be associated with proteins involved in cytoskeletal remodeling, cell adhesion and the recruitment of components of the translation machinery. FRS2’s integration into lipid rafts [64] could explain its distinct localization pattern, as these membrane microdomains have been reported to accumulate at the same subcellular compartments [65].

Oncogenic transformation through FRS2 amplifications and overexpression has been documented in several cancers [10–14], where higher expression levels of FRS2 correlate with increased anchorage-independent growth, proliferation and tumor formation *in vivo* [11,13]. Collectively, this data supports a role of FRS2 in promoting aggressive cellular phenotypes and identifies FRS2 as a potential target in MB and other tumors driven by activated FGFR or increased FRS2 expression [9,35,66–68]. Target engagement and repression of FRS2 functions by small molecule compounds [35] or pharmacological inhibition of FRS2 membrane association [67] indicate potential routes for therapeutic intervention. Redox regulation was recently found to subtly modulate the binding properties of the FRS2 PTB domain, offering a potential space for therapeutic interventions [69]. Our data provide interaction candidates to explore further for their context-dependent contribution of FGFR signaling and are consistent with a model in which modifications of the proximal FRS2 interactome could influence the selective regulation of its tumor-promoting activities. Additional studies are warranted to elucidate how the spatial and structural regulation of the FRS2 interactome imparts FGFR-driven oncogenic processes, ultimately to inform the development of next-generation therapeutics that more effectively and selectively disrupt pathological FGFR signaling.

## Conclusions

In summary, our findings position FRS2 as a protein associated with linking FGFR signaling to cytoskeletal remodeling, cell adhesion, and protein synthesis. Its overexpression in MB tumors and association with more aggressive, metastatic phenotypes highlight its potential clinical relevance. Therapeutic strategies aimed at selectively perturbing the FRS2 interactome hold promise for disrupting FGFR-driven migration and invasion in malignant tumors.

## Declarations

### Ethics approval and consent to participate

Not applicable

### Consent for publication

All authors have approved the submission of the manuscript

### Availability of data and materials

The acquired MS data are available on the ProteomeXchange Consortium via the PRIDE (http://www.ebi.ac.uk/pride) partner repository. See Material and Methods section for the respective identifiers. All other data that supports the findings of this study are available from the corresponding author on reasonable request.

### Competing interests

The authors declare that they have no conflict of interest.

### Funding

This study was funded by SNF-Sinergia project CRSII5_202245/1 and by Childhood Cancer Research Foundation Switzerland.

### Authors’ contributions

L.L.K.: Conceptualization, investigation, methodology, formal analysis, visualization, writing: Original draft preparation. D.V.: Methodology, investigation. B.C.: Methodology, data curation, formal analysis. M.T.S.: Methodology, investigation. D.H.: Investigation. M-S.L.: Methodology, investigation. M.B.: Conceptualization, supervision, formal analysis, data curation, funding acquisition, project administration, writing: Review & editing.

## Acknowledgments

We thank Guillaume Jacquemet from the University of Turku for his help and advice regarding the APEX2-based proximity labeling assay. Further thanks go to Shen Yan, who performed the flow cytometry analysis of the UW228 FRS2-NG^high^ cells. Mass-spectrometry was performed at the Functional Genomic Center Zürich. Special thanks go to Jonas Grossmann, Tobias Kockmann and Antje Dittmann for their support in planning and analyzing the mass-spectrometry assays. Some parts of Figure 4A were created with Created with BioRender.com.

---

**Figure S1**

**(A)** Kaplan–Meier survival probability curves for PanCancer patients with elevated (red) or unaltered (blue) *FRS2* mRNA levels. **(B)** Absolute number of TCGA PanCancer Atlas patients harboring altered *FRS3* mRNA levels, stratified by diagnosis. **(C)** Relative frequency of elevated *FRS3* mRNA levels per tumor diagnosis. **(D)** Kaplan–Meier survival probability curves for PanCancer patients with elevated (red) or unaltered (blue) *FRS3* mRNA levels. **(E)** Proportions of *FRS2* genomic alteration types stratified by tumor diagnosis. **(F)** Comparison of *FRS2* and *FRS3* mRNA expression z-scores in normal cerebellum (CB) versus medulloblastoma (MB) tumor samples from the GSE124814 dataset. Statistical significance was assessed by unpaired two-tailed Mann–Whitney test: ns p > 0.05, *p < 0.05, ***p < 0.0001. **(G)** *FRS2* and *FRS3* mRNA abundance in relation to metastatic status in the GSE124814 dataset (restricted to patients with available metastasis annotation). M0: no metastases detected; M+: metastases detected.

**Figure S2**

**(A)** IB analysis comparing FRS2 phosphorylation and MAPK/ERK and AKT pathway activation in wild-type and FRS2-mNG-expressing UW228 cells. **(B)** Proliferation competition assay using co-cultured wild-type and UW228 FRS2-mNG/mCh-Nuc UW228 cells. The relative proportion of FRS2-mNG-positive cells was assessed by flow cytometry using the mNeonGreen channel. **(C, D)** Formal hypothesis testing and effect size matrices from live-cell tracking, comparing mean speed (C) and directionality (D) of UW228 cells overexpressing LA-EGFP or FRS2-mNG (FRS2−: mixed FRS2-mNG expression; FRS2+: high FRS2-mNG expression; LA-EGFP: control). **(E)** Spheroid invasion assay (SIA) comparing wild-type and FRS2-mNG-overexpressing UW228 cells in the presence of 100 ng/mL bFGF. Dots represent individual spheroids (each comprising >2,500 cells) from technical replicates of n = 1 experiment. Statistical significance was assessed by one-way ANOVA with Kruskal–Wallis and multiple comparisons correction: *p ≤ 0.05, **p ≤ 0.01, ***p ≤ 0.001, ****p ≤ 0.0001. **(F)** Validation of FRS2 knockdown efficiency for siFRS2 #2 and siFRS2 #3 by RT-qPCR in wild-type ONS-76 cells. **(G)** Immunofluorescence analysis of endogenous FRS2 localization in wild-type UW228 cells.

**Figure S3**

**(A)** Determination of the optimal bFGF concentration for FGFR signaling activation in D425 cells by IB analysis following 5 min stimulation. **(B)** Total proteome changes in D425 cells stimulated with 50 ng/mL bFGF for 5 min. Gene names of significantly upregulated proteins (FDR ≤ 0.1, log₂FC ≥ 0.6) are labeled in the upper right quadrant.

**Figure S4**

**(A)** Subcellular localization of FRS2-APEX2-V5 in UW228 cells assessed by IF analysis. The transgene was detected using an anti-V5 antibody. **(B)** Comparative IB analysis of FRS2 phosphorylation in wild-type UW228 and UW228 FRS2-APEX2-V5 (F-APX) cells following bFGF stimulation (50 ng/mL, 5 min). **(C)** IB analysis of eluates from wild-type UW228 (WT) and UW228 F-APX pull-downs using an FGFR peptide as bait. **(D)** IB detection of biotinylated proteins using streptavidin-HRP, comparing samples with and without pellet snap-freezing prior to lysis. **(E)** Comparison of filtration versus acetone precipitation for removal of excess biotin prior to pull-down, and their respective effects on biotinylated protein recovery. **(F)** Subcellular localization distribution of bFGF-induced FRS2 proximal interactor candidates identified by APEX2-based proximity labeling. **(G)** Overlap between APEX2-labeled proteins identified in the FRS2 proximal proteome under serum-free and bFGF-stimulated conditions combined and previously reported FRS2 interactors catalogued in the BioGRID database.

**Figure S5**

**(A)** Validation of APEX2-labeled proteins TJP1, WASF2, YAP1, and EZR. Immunoblot analysis following streptavidin-HRP pulldown, probed with antibodies against the proteins indicated to the right of each panel. **(B)** Venn diagram visualizing shared hits across the following datasets: phosphoproteome analysis, APEX2 proximity labeling, co-immunoprecipitation, and predicted FRS2 interactors from the BioGRID database. **(C–E)** Biotinylated peptides identified by APEX2 proximity labeling (highlighted in red) mapped onto the predicted AlphaFold structures of WASF2 (C), PDLIM1 (D), and TJP1 (E). Schematic representations of the folded domain architecture are shown below each structure, with the general molecular functions of individual domains indicated.

**Figure S6**

**(A)** RT-qPCR analysis of *FRS2*, *FRS3*, and *FGFR1–4* mRNA levels in wild-type HAP1 and HAP1 FRS2-KO cells. Statistical significance was assessed by unpaired two-tailed Student’s t-test: ns p > 0.05, ***p ≤ 0.001. **(B)** RT-qPCR analysis of *FRS2* and *FRS3* mRNA expression in wild-type HAP1 cells. **(C)** Magnified views of IFA images from Figure 6A, showing wild-type HAP1 and HAP1 FRS2-KO cells stimulated with 50 ng/mL bFGF for 1 h. F-actin is shown in green (phalloidin) and TJP1 in red. **(D)** IB analysis of FRS2 interactor expression in wild-type HAP1 and HAP1 FRS2-KO cells.

**Figure S7**

**(A, B)** Z-scores of *WASF2* (A) and *PDLIM1* (B) mRNA expression across two control datasets (normal brain regions and cerebellum, shown in green) and six primary MB patient datasets (shown in blue), analyzed on the Affymetrix U133P2 platform. Significant differences relative to the healthy cerebellum control were assessed by one-way ANOVA with Kruskal–Wallis and Dunn’s multiple comparisons tests: ns p ≥ 0.05, *p ≤ 0.0332, **p ≤ 0.0021, ***p ≤ 0.0002, ****p ≤ 0.0001. **(C, D)** Left: Z-scores of *PDLIM1* (C) and *WASF2* (D) mRNA expression in cerebellar control (n = 109) and MB tumor samples (n = 1’314). Right: Z-scores of *PDLIM1* and *WASF2* expression in non-metastatic MB patients (M0) versus patients with tumor cells detectable in the cerebrospinal fluid (M+; GSE124814 dataset). Statistical significance was assessed by unpaired two-tailed Mann–Whitney test: ns p ≥ 0.05, ****p ≤ 0.0001. **(E, F)** Kaplan–Meier curves of overall survival in MB patients stratified by high (red) or low (blue) tumor expression of *PDLIM1* (E) and *WASF2* (F). **(G)** Left: Correlation plots of *FRS2* mRNA expression z-scores versus *PDLIM1* or *WASF2* across MB subgroups. Right: Corresponding slope statistics.

## Notes

http://www.ebi.ac.uk/pride

### Competing Interest Statement

The authors have declared no competing interest.

### Summary of Updates

Title adjusted to remove hub terminology Additional data and quantifications were added and text rephrased to avoid overstatements.

