## Supplementary material for "Fibroblast growth factor receptor substrate 2 interactome mapping reveals novel candidate interactors associated with migration and invasion": SF1

**Supplementary methods file:**

**Table 1**: Used cell lines

| Name | Origin | Authentication/testing | Medium |
| --- | --- | --- | --- |
| UW228 | John Silber, Seattle, USA | Single Nucleotide Polymorphism (SNP) typing, 02/2022  Regular LookOut^®^ Mycoplasma PCR Detection kit (Sigma Aldrich, #MP0035) | DMEM medium (Gibco, #11965092) supplemented with 1% v/v GlutaMax (Gibco, #35050-038), 10% v/v FBS (Sigma, #S0615), 1% v/v sodium pyruvate (ThermoFisher, #11360-070) and 1% v/v penicillin/streptomycin (Gibco, #15140-122) |
| ONS-76 | Michael Taylor (SickKidds, Toronto, Canada) | Single Nucleotide Polymorphism (SNP) typing, 02/2022  Regular LookOut^®^ Mycoplasma PCR Detection kit (Sigma Aldrich, #MP0035) | RPMI 1640 (R0883 Sigma) supplemented with 10% v/v FBS (Sigma, #S0615) and 1% v/v penicillin/streptomycin (Gibco, #15140-122) |
| D425-Med (D425) | Henry Friedman lab, Duke Unviersity, USA | Single Nucleotide Polymorphism (SNP) typing, 02/2022  Regular LookOut^®^ Mycoplasma PCR Detection kit (Sigma Aldrich, #MP0035) | IMEM medium (Gibco, #10373-017) supplemented with 10% v/v FBS (Sigma, #S0615) and 1% v/v penicillin/streptomycin (Gibco, #15140-122) |
| HEK Lenti-X 293 | Beat Schäfer, UZH, Switzerland | Regular LookOut^®^ Mycoplasma PCR Detection kit (Sigma Aldrich, #MP0035) | DMEM medium (Gibco, #11965092) supplemented with 1% v/v GlutaMax (Gibco, #35050-038), 10% v/v FBS (Sigma, #S0615), 1% v/v sodium pyruvate (ThermoFisher, #11360-070) and 1% v/v penicillin/streptomycin (Gibco, #15140-122) |

**Table 2**: Primary antibodies used in Western Blot

| Antibody | Source | Dilution | Company | Catalog Number |
| --- | --- | --- | --- | --- |
| FRS2 mAb | Rabbit | 1:1000 | Abcam | ab183492 |
| FRS2 pAb | Rabbit | 1:1000 | Abcam | ab137458 |
| FRS2 | Mouse | 1:1000 | Merck | 05-502 |
| pFRS2 (Y426) | Rabbit | 1:1000 | Cell Signaling Technology | 3861 |
| ERK 1/2 | Rabbit | 1:1000 | Cell Signaling Technology | 9102 |
| pERK 1/2 (T202/Y204) | Rabbit | 1:1000 | Cell Signaling Technology | 9101 |
| AKT | Rabbit | 1:1000 | Cell Signaling Technology | 9272 |
| pAKT (S473) | Rabbit | 1:1000 | Cell Signaling Technology | 4060 |
| GAP1 | Rabbit | 1:1000 | Cell Signaling Technology | 3232 |
| pGAP1 (Y627) | Rabbit | 1:1000 | Cell Signaling Technology | 3233 |
| SHP2 | Rabbit | 1:1000 | Cell Signaling Technology | 3397 |
| pSHP2 (Y580) | Rabbit | 1:1000 | Cell Signaling Technology | 3703 |
| pSHP2 (Y542) | Rabbit | 1:1000 | Cell Signaling Technology | 3751 |
| TJP1/ZO-1 | Rabbit | 1:1000 | Cell Signaling Technology | 8193 |
| WASF2/  WAVE-2 | Rabbit | 1:1000 | Cell Signaling Technology | 3659 |
| YAP | Rabbit | 1:1000 | Cell Signaling Technology | 14074 |
| Ezrin/Radixin/  Moesin | Rabbit | 1:1000 | Cell Signaling Technology | 3142 |
| PDLIM1 | Mouse | 1:1000 | Abcam | ab129015 |
| Streptavidin-HRP | - | 1:1000 | Cell Signaling Technology | 3999 |
| V5-tag | Rabbit | 1:1000 | Cell Signaling Technology | 13202 |
| Histone H3 | Rabbit | 1:1000 | Cell Signaling Technology | 4499 |
| GAPDH | Rabbit | 1:1000 | Cell Signaling Technology | 2118 |

**Table 3**: Primary antibodies used in IFA

| Antibody | Source | Dilution | Company | Catalog Number |
| --- | --- | --- | --- | --- |
| FRS2 | Rabbit | 1:100 | Santa Cruz | sc-8318 |
| V5-Tag | Rabbit | 1: 500 | Cell Signaling Technology | 13202 |

**Table 4**: Secondary antibodies used in Western Blot

| Antibody | Source | Dilution | Company | Catalog Number |
| --- | --- | --- | --- | --- |
| Anti-mouse IgG, HRP-linked | Horse | 1:5000 | Cell Signaling Technology | 7076 |
| Anti-rabbit IgG, HRP-linked | Goat | 1:5000 | Cell Signaling Technology | 7074 |

**Table 5**: Secondary antibodies used in IFA

| Antibody | Source | Dilution | Company | Catalog Number |
| --- | --- | --- | --- | --- |
| Alexa Fluor 647 – Anti-rabbit IgG (H+L) | Donkey | 1:250 | Jackson ImmunoResearch | 711-605-152 |

Primary and secondary antibodies used in Western Blot and IFA were diluted with 1× TBST-T with 5% m/v non-fat dry milk (WB total protein) or 5% m/v bovine serum albumin (WB phospho-protein) and 5% FBS in PBS (IFA).

**Table 6**: Lentiviral transgene vectors

| Vector Name | Vector ID | Company | Selection Marker |
| --- | --- | --- | --- |
| pLV[Exp]-Bsd-CMV>  {hFRS2_mNeonGreen}/3xGS | VB220316-1595dwh | VectorBuilder | Bsd (blasticidin resistance gene |
| pLV[Exp]-Bsd-CMV>  hFRS2[NM_001278357.2]  (ns)/3xGS:{APEX}/3xGS/V5 | VB221102-1154pbp | VectorBuilder | Bsd (blasticidin resistance gene |

Transgene coding sequences in lentiviral vectors:

**pLV_Bsd-CMV_hFRS2_3xGS_mNeonGreen**

MGSCCSCPDKDTVPDNHRNKFKVINVDDDGNELGSGIMELTDTELILYTRKRDSVKWHYLCLRRYGYDSNLFSFESGRRCQTGQGIFAFKCARAEELFNMLQEIMQNNSINVVEEPVVERNNHQTELEVPRTPRTPTTPGFAAQNLPNGYPRYPSFGDASSHPSSRHPSVGSARLPSVGEESTHPLLVAEEQVHTYVNTTGVQEERKNRTSVHVPLEARVSNAESSTPKEEPSSIEDRDPQILLEPEGVKFVLGPTPVQKQLMEKEKLEQLGRDQVSGSGANNTEWDTGYDSDERRDAPSVNKLVYENINGLSIPSASGVRRGRLTSTSTSDTQNINNSAQRRTALLNYENLPSLPPVWEARKLSRDEDDNLGPKTPSLNGYHNNLDPMHNYVNTENVTVPASAHKIEYSRRRDCTPTVFNFDIRRPSLEHRQLNYIQVDLEGGSDSDNPQTPKTPTTPLPQTPTRRTELYAVIDIERTAAMSNLQKALPRDDGTSRKTRHNSTDLPMGSGSGSVSKGEEDNMASLPATHELHIFGSINGVDFDMVGQGTGNPNDGYEELNLKSTKGDLQFSPWILVPHIGYGFHQYLPYPDGMSPFQAAMVDGSGYQVHRTMQFEDGASLTVNYRYTYEGSHIKGEAQVKGTGFPADGPVMTNSLTAADWCRSKKTYPNDKTIISTFKWSYTTGNGKRYRSTARTTYTFAKPMAANYLKNQPMYVFRKTELKHSKTELNFKEWQKAFTDVMGMDELYK-

**pLV_Bsd-CMV_hFRS2_3xGS_APEX2_3xGS_V5**

MGSCCSCPDKDTVPDNHRNKFKVINVDDDGNELGSGIMELTDTELILYTRKRDSVKWHYLCLRRYGYDSNLFSFESGRRCQTGQGIFAFKCARAEELFNMLQEIMQNNSINVVEEPVVERNNHQTELEVPRTPRTPTTPGFAAQNLPNGYPRYPSFGDASSHPSSRHPSVGSARLPSVGEESTHPLLVAEEQVHTYVNTTGVQEERKNRTSVHVPLEARVSNAESSTPKEEPSSIEDRDPQILLEPEGVKFVLGPTPVQKQLMEKEKLEQLGRDQVSGSGANNTEWDTGYDSDERRDAPSVNKLVYENINGLSIPSASGVRRGRLTSTSTSDTQNINNSAQRRTALLNYENLPSLPPVWEARKLSRDEDDNLGPKTPSLNGYHNNLDPMHNYVNTENVTVPASAHKIEYSRRRDCTPTVFNFDIRRPSLEHRQLNYIQVDLEGGSDSDNPQTPKTPTTPLPQTPTRRTELYAVIDIERTAAMSNLQKALPRDDGTSRKTRHNSTDLPMGSGSGSGKSYPTVSADYQDAVEKAKKKLRGFIAEKRCAPLMLRLAFHSAGTFDKGTKTGGPFGTIKHPAELAHSANNGLDIAVRLLEPLKAEFPILSYADFYQLAGVVAVEVTGGPKVPFHPGREDKPEPPPEGRLPDPTKGSDHLRDVFGKAMGLTDQDIVALSGGHTIGAAHKERSGFEGPWTSNPLIFDNSYFTELLSGEKEGLLQLPSDKALLSDPVFRPLVDKYAADEDAFFADYAEAHQKLSELGFADALQGSGSGSGKPIPNPLLGLDST-

**Table 7**: siRNA’s used

| **siRNA** | | | |
| --- | --- | --- | --- |
| **Name** | **Target Sequence** | **Company** | **Identifier** |
| siCTRL |  | Invitrogen, Silencer Select | 4390843 |
| siFRS2_2 | GGCUAUGACUCGAAUCUCU | Invitrogen, Silencer Select | s21261 |
| siFRS2_3 | GCAUAAUGGAACUUACAGA | Invitrogen, Silencer Select | s21262 |
