## Supplementary material for "Fibroblast growth factor receptor substrate 2 interactome mapping reveals novel candidate interactors associated with migration and invasion": SF10

### Figure 3A

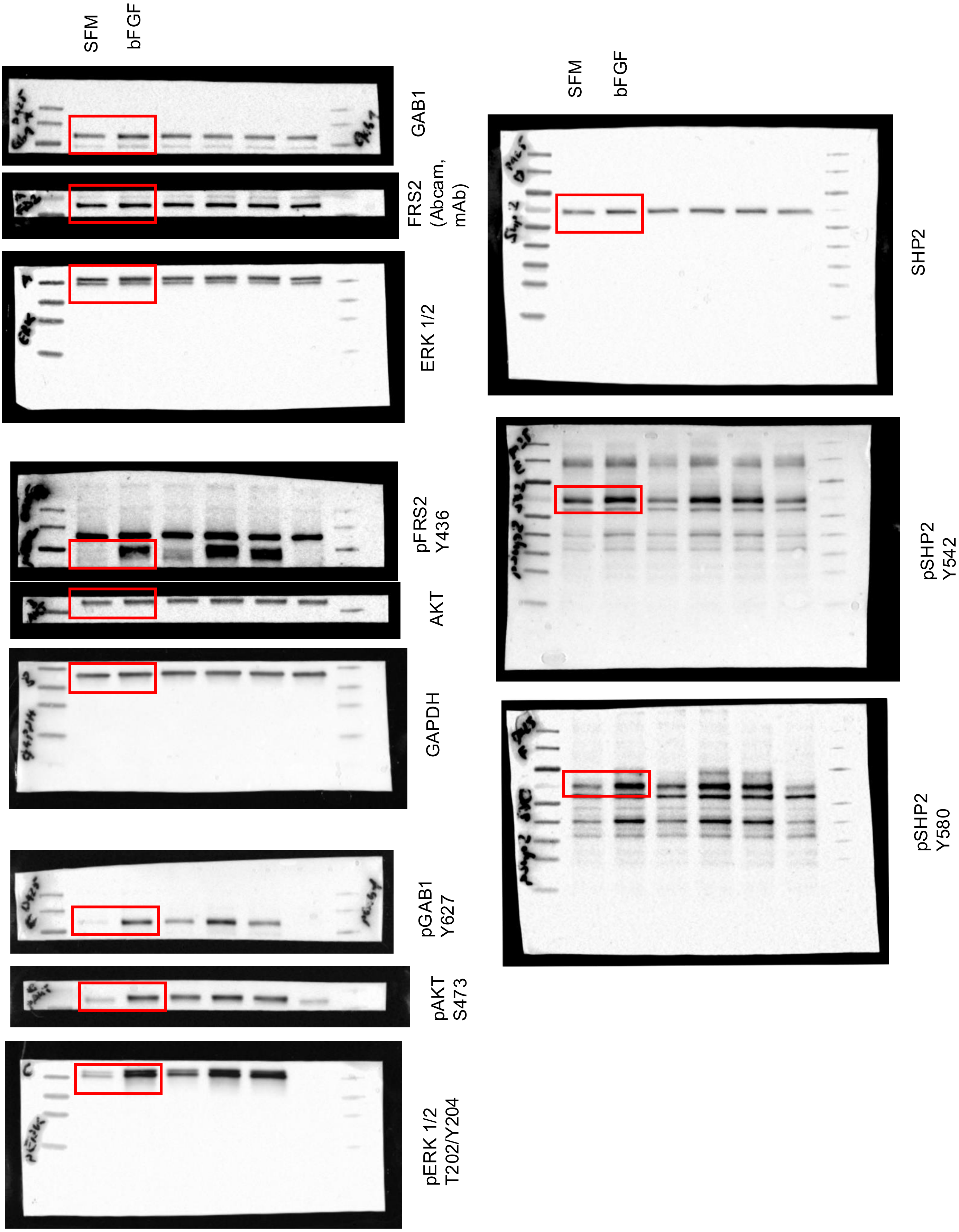
SFM
bFGF
GAB1
FRS2 (Abcam, mAb)
ERK 1/2
pFRS2 Y436
AKT
GAPDH
pGAB1 Y627
pAKT S473
pERK 1/2 T202/Y204
SFM
bFGF
SHP2
pSHP2 Y542
pSHP2 Y580

### Figure 3E

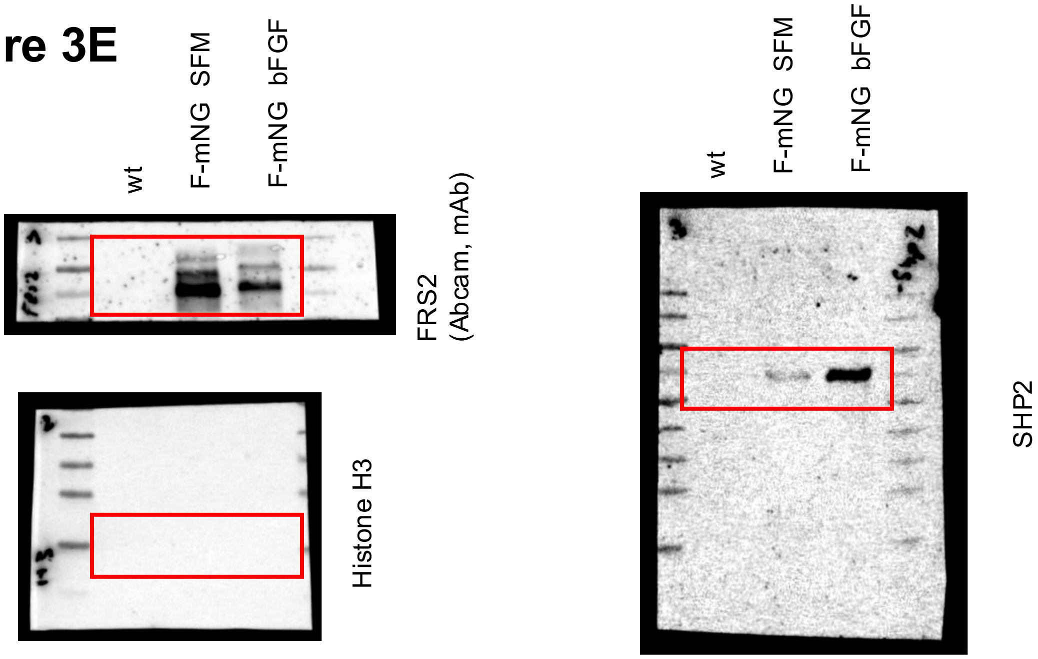


### Figure 4B

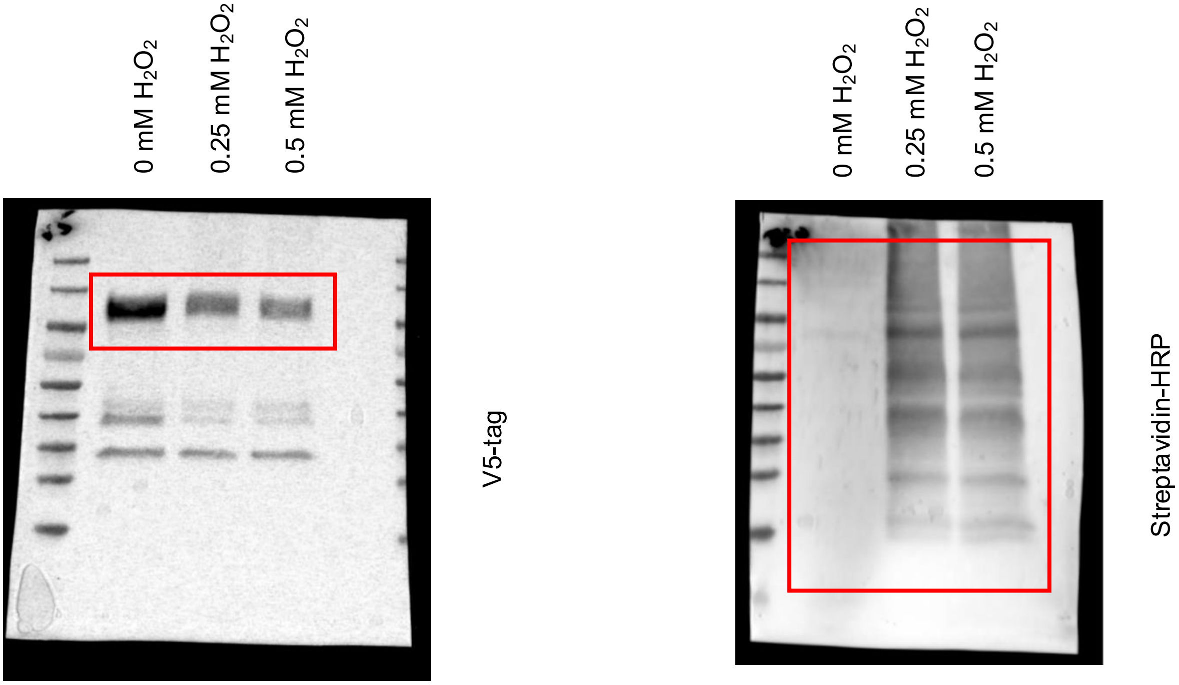

### Figure 4C

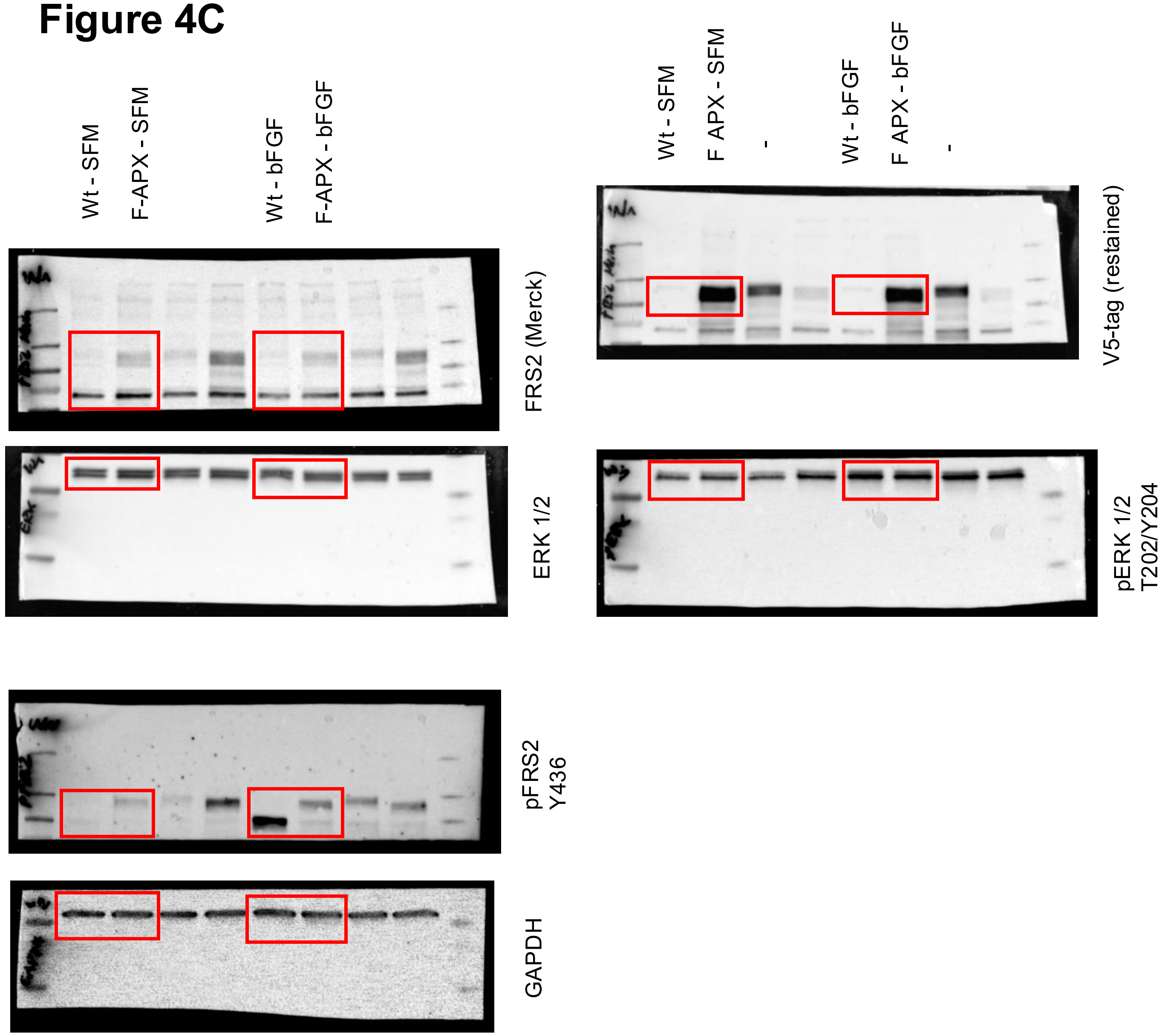

Wt - SFM
F-APX - SFM
Wt - bFGF
F-APX - bFGF
FRS2 (Merck)
ERK 1/2
pFRS2
Y436
GAPDH
Wt - SFM
F APX - SFM
-
Wt - bFGF
F APX - bFGF
-
V5-tag (restained)
pERK 1/2
T202/Y204

### Figure 5B

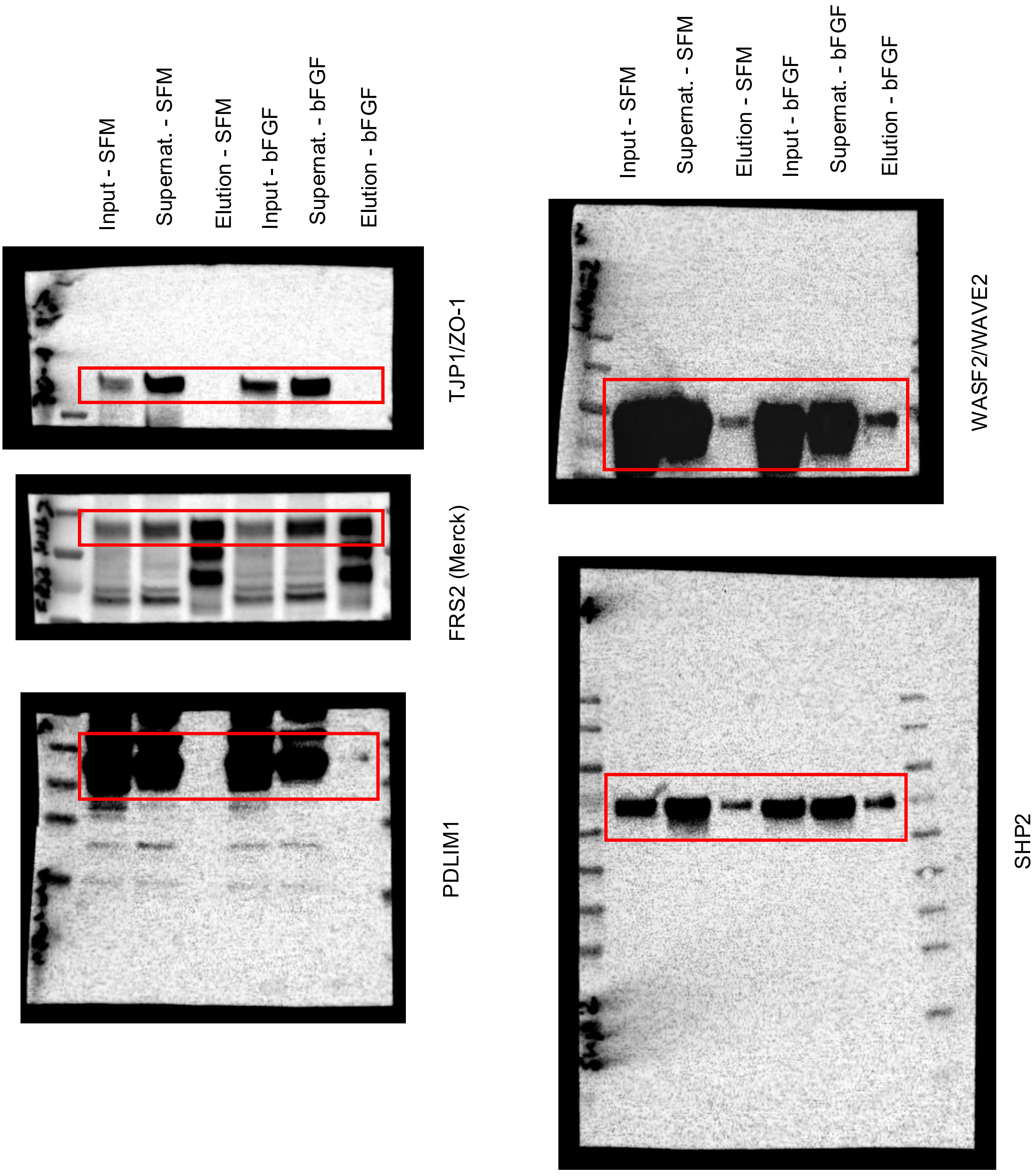

Input - SFM
Supernat. - SFM
Elution - SFM
Input - bFGF
Supernat. - bFGF
Elution - bFGF
TJP1/ZO-1
FRS2 (Merck)
PDLIM1
WASF2/WAVE2
SHP2

### Figure S2A

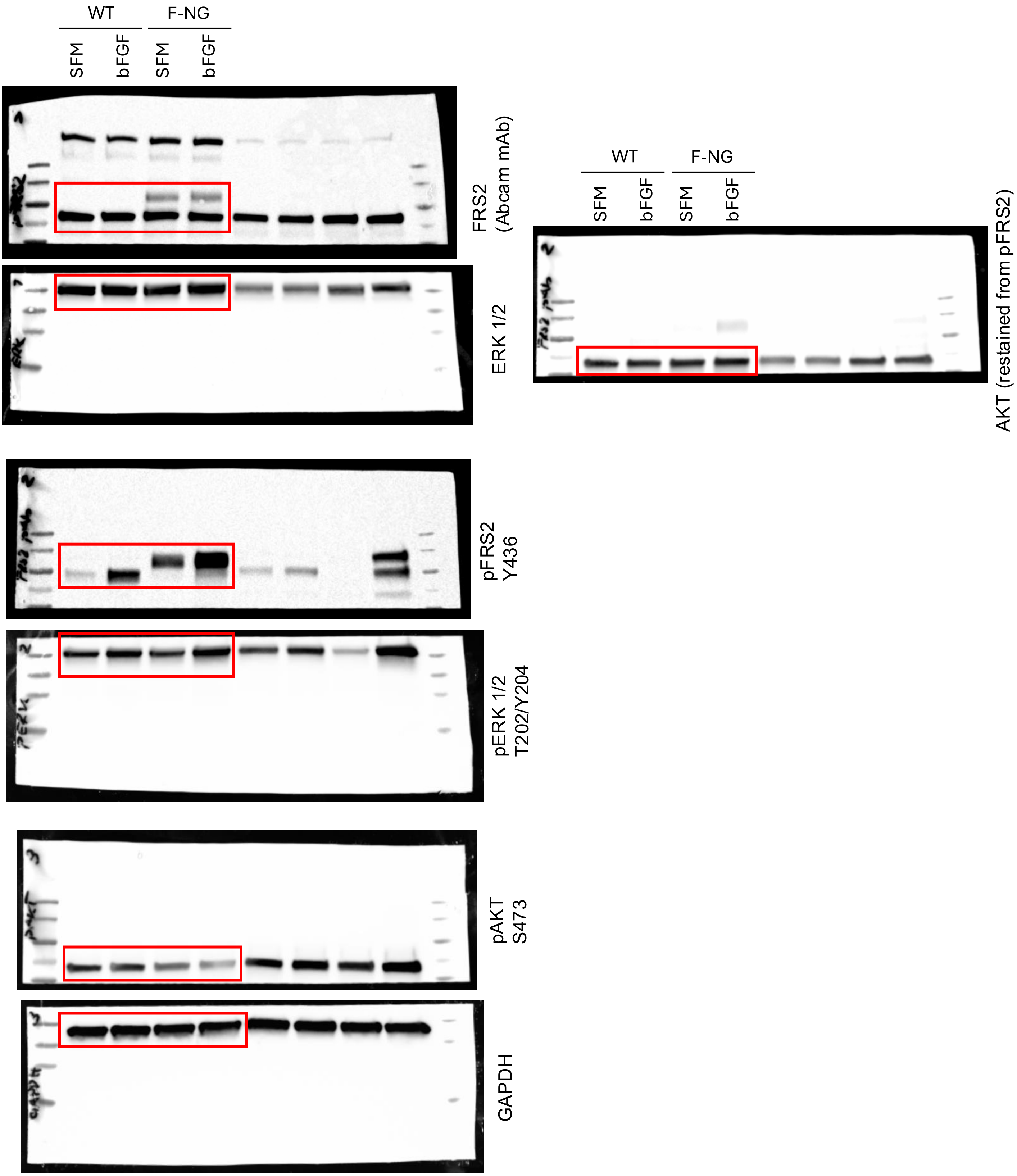

WT
F-NG
SFM
bFGF
SFM
bFGF
FRS2
(Abcam mAb)
ERK 1/2
pFRS2
Y436
pERK 1/2
T202/Y204
pAKT
S473
GAPDH
WT
F-NG
SFM
bFGF
SFM
bFGF
AKT (restained from pFRS2)

### Figure S3A

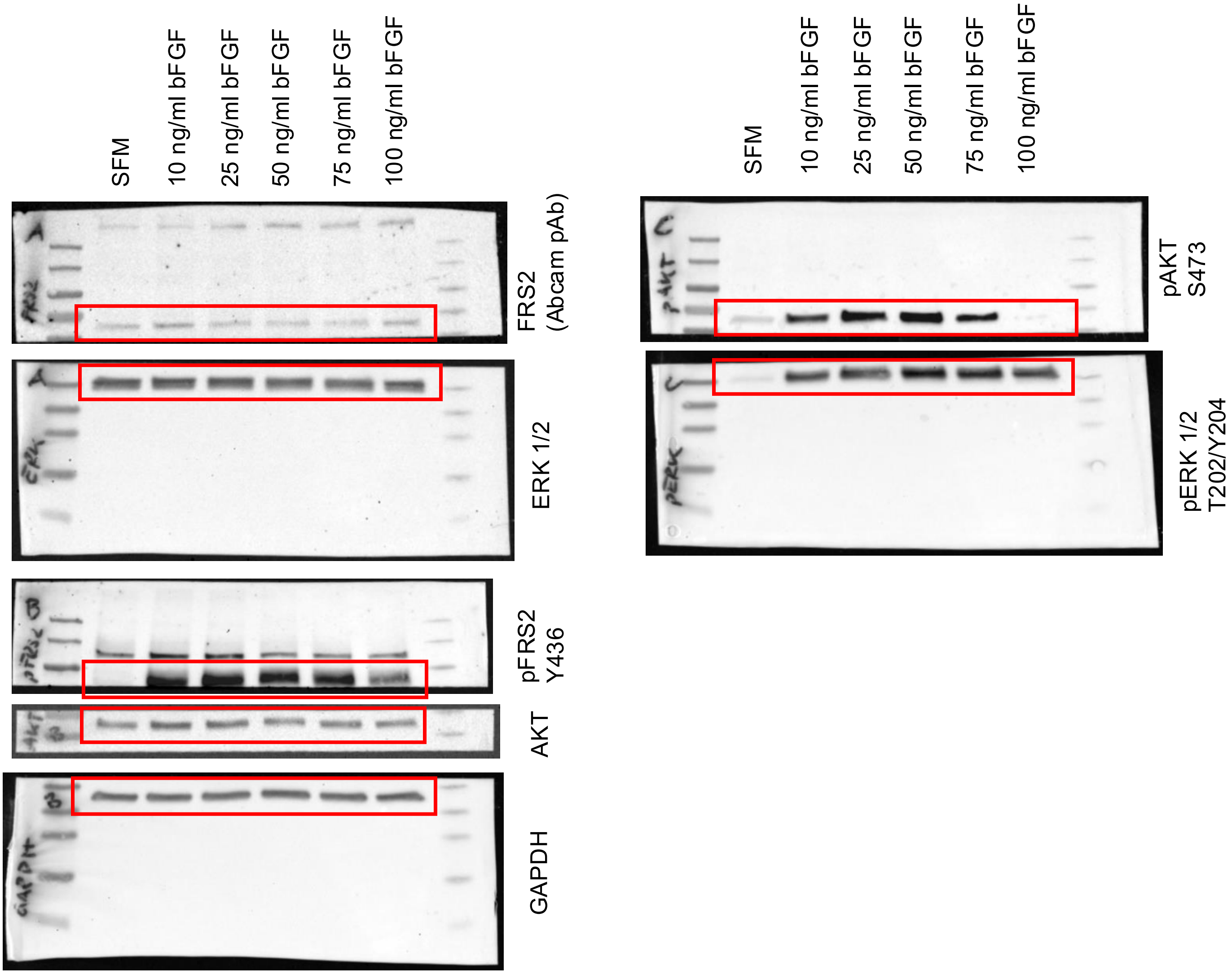
SFM
10 ng/ml bFGF
25 ng/ml bFGF
50 ng/ml bFGF
75 ng/ml bFGF
100 ng/ml bFGF
FRS2 (Abcam pAb)
ERK 1/2
pFRS2 Y436
AKT
GAPDH
SFM
10 ng/ml bFGF
25 ng/ml bFGF
50 ng/ml bFGF
75 ng/ml bFGF
100 ng/ml bFGF
pAKT S473
pERK 1/2 T202/Y204

### Figure S4A

Wt - SFM
Wt - bFGF
F-APX - SFM
F-APX - bFGF

Wt - SFM
Wt - bFGF
F-APX - SFM
F-APX - bFGF

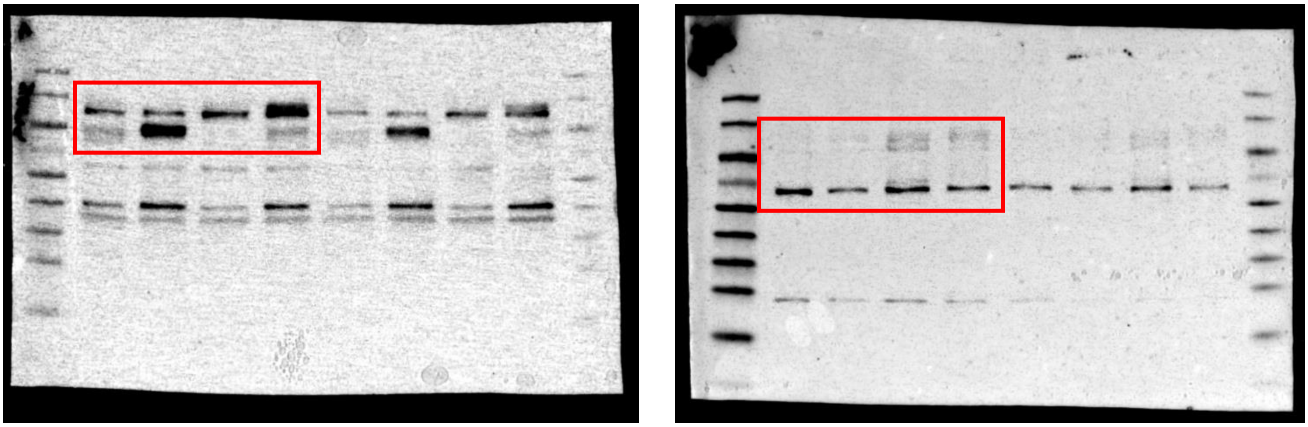

pFRS2
Y436

FRS2 (Merck)

### Figure S4C

WT
F-APX

WT
F-APX

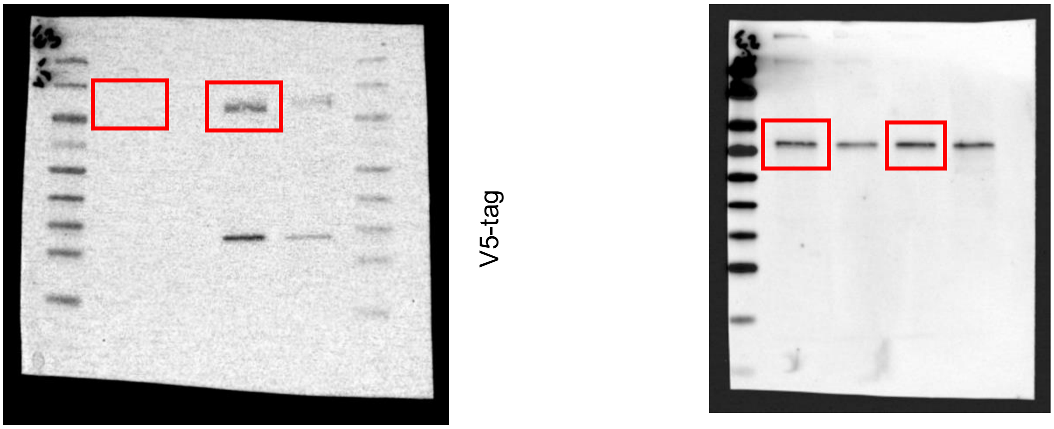

V5-tag

FRS2 (Abcam, mAb)

**Figure S4D**

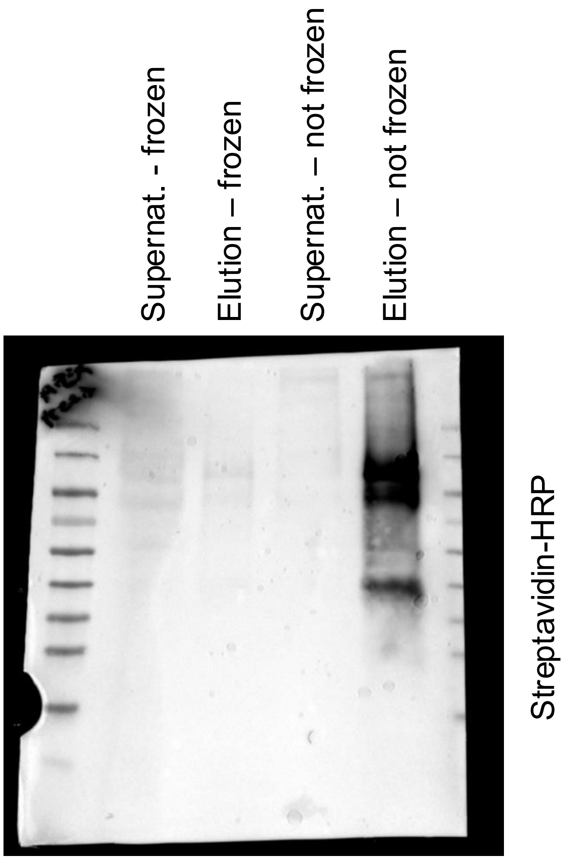


**Figure S4E**

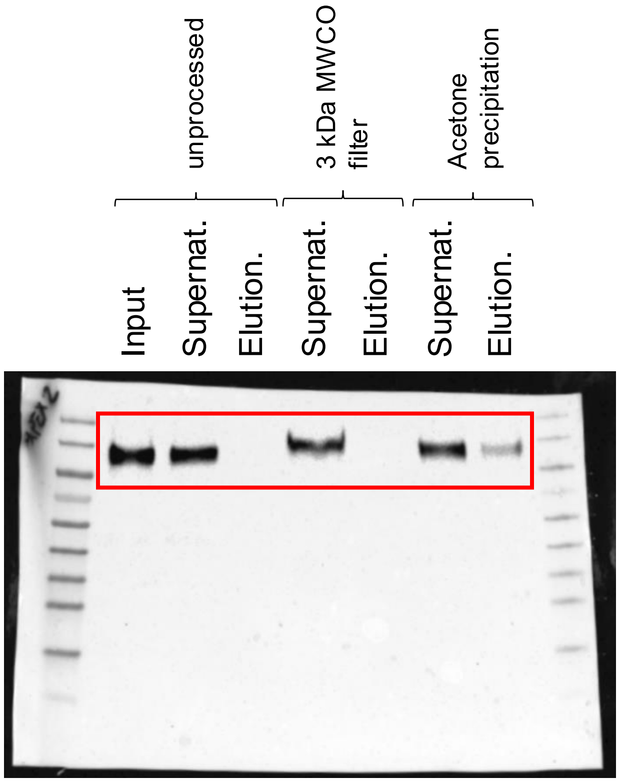


V5-tag

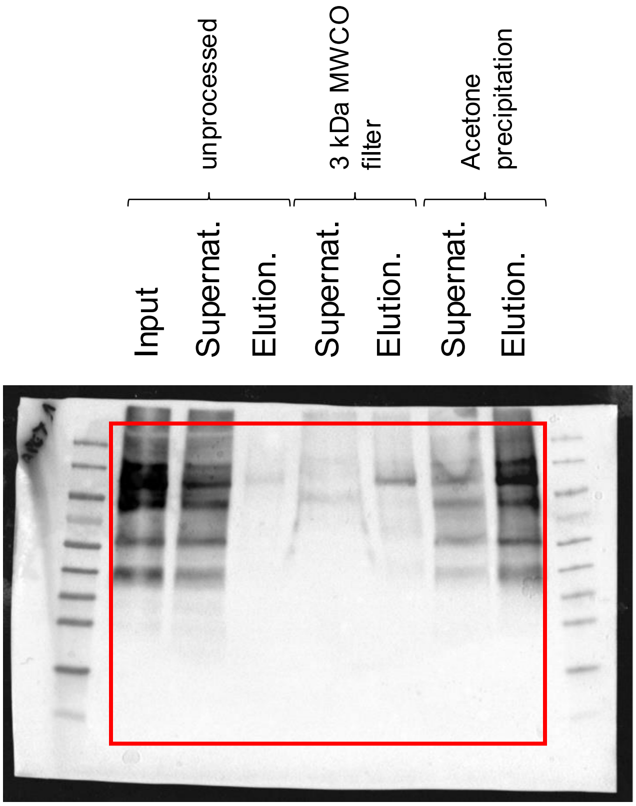


Streptavidin-HRP

### Figure S5A

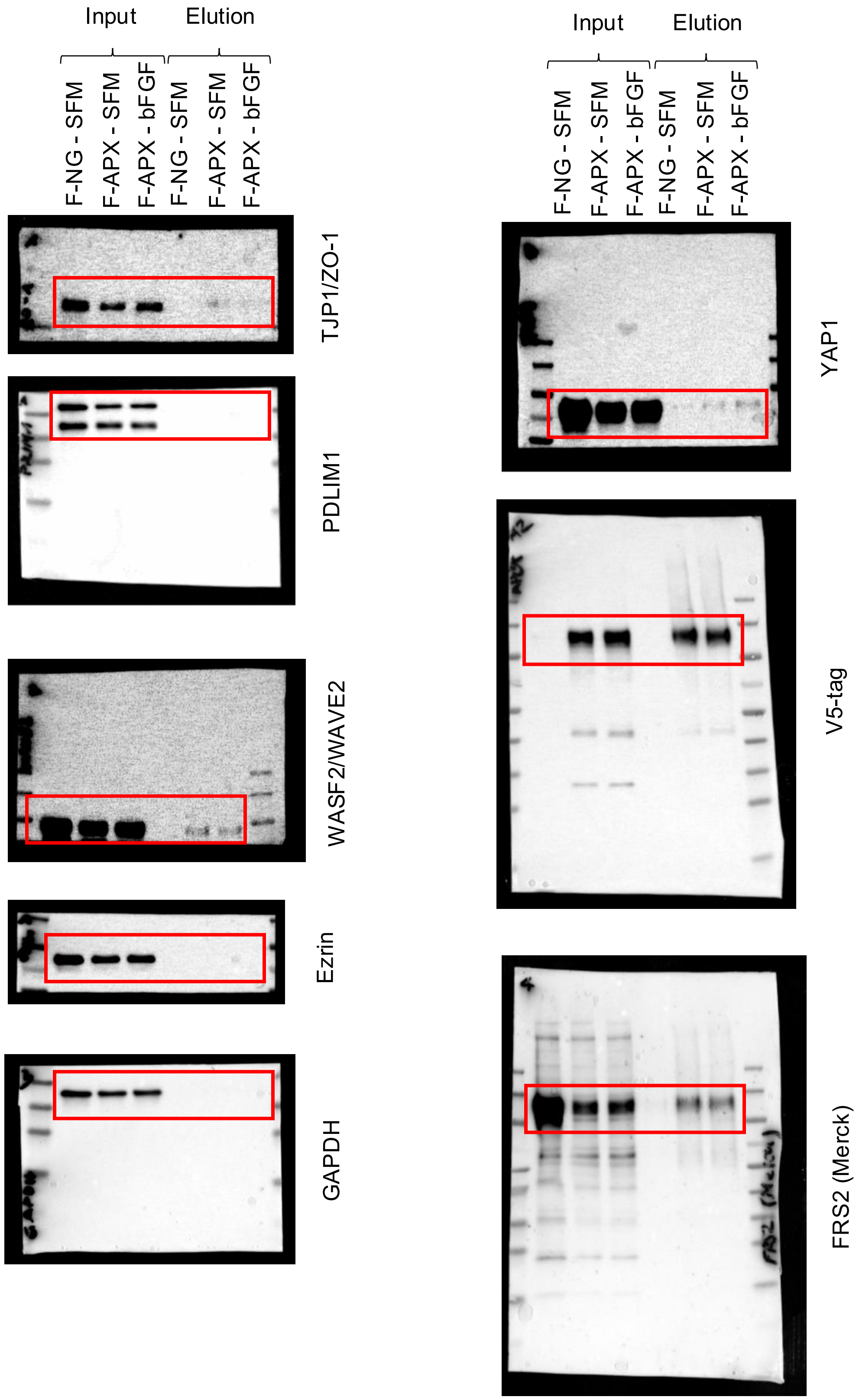
Input
Elution
F-NG - SFM
F-APX - SFM
F-APX - bFGF
F-NG - SFM
F-APX - SFM
F-APX - bFGF
TJP1/ZO-1
PDLIM1
WASF2/WAVE2
Ezrin
GAPDH
Input
Elution
F-NG - SFM
F-APX - SFM
F-APX - bFGF
F-NG - SFM
F-APX - SFM
F-APX - bFGF
YAP1
V5-tag
FRS2 (Merck)

### Figure S6C

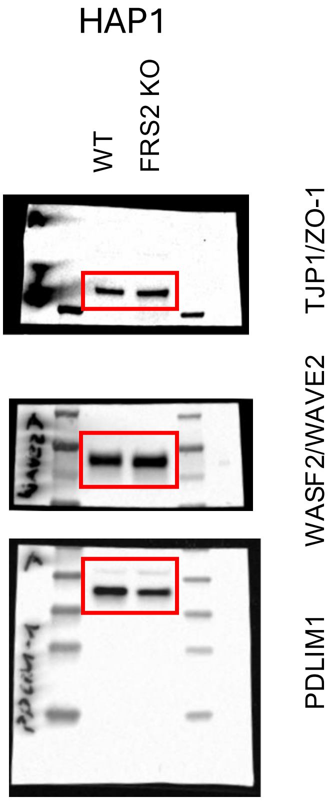

HAP1
WT
FRS2 KO
TJP1/ZO-1
WASF2/WAVE2
PDLIM1


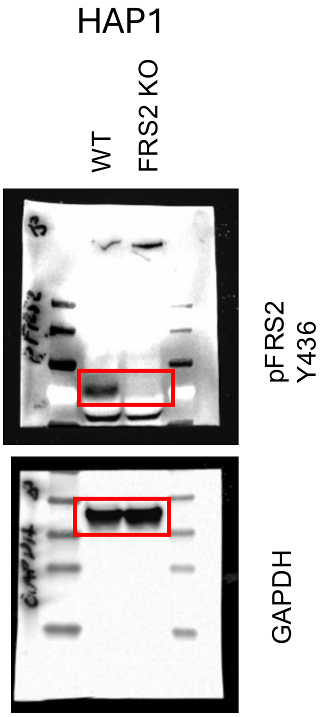

HAP1
WT
FRS2 KO
pFRS2 Y436
GAPDH
